# Predicting Head and Neck Squamous Cell Carcinoma outcomes using long-term Patient-Derived Tumor Organoids

**DOI:** 10.64898/2026.03.22.713356

**Authors:** Marion Perréard, Jordane Divoux, Félicie Perrin, Romane Florent, Lucie Lecouflet, Guillaume Desmartin, Lucie Thorel, Florence Giffard, Sarah Burton, Jade Richard, Jean-Michel Grellard, Esther Lebreton, Emilie Brotin, Céline Villenet, Shéhérazade Sebda, Jean-Pascal Meneboo, Anamika Pandey, Valentin Harter, Corinne Jeanne, Céline Bazille, Audrey Lasne-Cardon, Maxime Humbert, Gaurav Kumar Pandey, Vianney Bastit, François Christy, Juliette Thariat, Nicolas Vigneron, Emmanuel Babin, Martin Figeac, Matthieu Meryet-Figuière, Laurent Poulain, Louis-Bastien Weiswald

## Abstract

Head and neck squamous cell carcinoma (HNSCC) remains associated with substantial morbidity and a 5-year overall survival rate of approximately 60%, reflecting persistent radio- and chemo-resistance and the lack of effective precision medicine strategies. Patient-Derived Tumor Organoids (PDTO) constitute promising functional models that may predict individual treatment response. In this study, we generated PDTO from surgically resected HNSCC of the oral cavity, oropharynx, larynx, and hypopharynx. A total of 20 long-term PDTO lines were established, maintaining growth over seven passages and successfully cryopreserved, capturing the molecular and clinical diversity of the patient cohort. These PDTO faithfully recapitulated histological features, major tumor marker expression, and the genomic and transcriptomic landscapes of their tumors of origin, with stability over time. Functional assays revealed heterogeneous responses to cisplatin and X-rays. Importantly, *in vitro* sensitivity of PDTO was associated with clinical outcome of patients at 24 months. Cisplatin response of PDTO predicted prognosis with 66.7% sensitivity and 100% specificity, while X-ray response showed 91.7% sensitivity and 75% specificity. Notably, all patients whose PDTO were classified as resistant to both cisplatin and X-rays experienced relapse and/or death within 24 months. Collectively, the successful long-term expansion and cryopreservation of HNSCC PDTO establish a stable and scalable preclinical resource that captures the molecular and clinical heterogeneity of the disease. This biobank provides a valuable platform for mechanistic studies and for the evaluation of innovative therapeutic strategies. This cohort represents one of the largest clinically annotated HNSCC PDTO collections to date, demonstrating a robust association between PDTO response to cisplatin and X-rays and patient prognosis. These findings support the predictive potential of PDTO-based functional assays and argue for their integration into standardized, rapid, and miniaturized precision oncology workflows for HNSCC.

## Introduction

Head and neck cancers (HNC) are the seventh most common cancer worldwide, accounting for approximately 890,000 new cases and 450,000 deaths in 2020 [1]. Head and Neck Squamous Cell Carcinoma (HNSCC) represents more than 90% of HNC and encompass a spectrum of heterogeneous malignancies originating from three major anatomical sites: the oral cavity, pharynx, and larynx, although some studies also classify sinonasal cavity as HNSCC [2]. Tobacco use and excessive alcohol consumption are considered among the most significant risk factors [3] as well as infection with human papillomavirus (HPV) contributing for HPV for around 22% of oropharyngeal cancer [4]. Despite advances in therapy, the 5-year survival rate for HNSCC has been only moderately improved over the past three decades [5] and remains at 60%, although HPV-positive HNSCC have a more favorable outcome, probably due to their increased sensitivity to chemoradiotherapy [6]. Currently available treatment options consist of surgery, radiation, chemotherapy, targeted therapy and immunotherapy, administered in single or multi-modality. Indeed, patients with locally advanced HNSCC received radiochemotherapy or undergo surgery followed by adjuvant radiotherapy alone or combined with systemic treatments such as cisplatin or cetuximab. However, these treatments are associated with significant morbidity and acquisition of chemo- or radio-resistance [7]. Therefore, it is crucial to implement precision medicine to tailor treatment, i.e. select the most effective one and avoid therapies that will not work or carry a high risk of toxicity.

To date, genomics has been the main tool in guiding precision medicine. This approach is based on the idea that somatic genetic alterations can be identified and matched with drugs targeting those abnormalities to improve treatment outcomes. However, studies have generally shown that most cancer patients who receive genomic testing do not benefit from a genomic precision medicine strategy [8], since many of them lack actionable alterations and matched treatments. This revealed the interest to explore additional approaches, such as functional precision medicine. This strategy relies on patient-derived tumor models, which are exposed to therapies and which response is measured to predict clinical response [8]. In this context, the recent advent of Patient-Derived Tumor Organoids (PDTO) offers promising opportunities to facilitate personalized treatment strategies. PDTO are grown from patient tumor cells in 3D extracellular matrix surrounded by culture media supplemented with a cocktail of growth factors and inhibitors of signaling pathways to mimic *in vivo* niche conditions [9]. They can be rapidly expanded and are representative of their parental tumor tissue. Above all, increasing evidence indicates that PDTO are able to recapitulate response of the patient, although most of clinical studies were based on small sample size [10]. PDTO cultures have already been successfully established from patients with HNSCC, although with variable success rate of establishment ranging from 28% to 93% [11–21]. Interestingly, six of these studies [12, 13, 15, 16, 19, 20] showed a significant correlation between PDTO response and clinical outcomes despite the small sample sizes, ranging from 3 to 14 patients.

Here, we established a panel of 20 long-term PDTO from patients with HNSCC and performed a comprehensive characterization of these models before assessing their potential to predict accurately the patients’ response to treatments.

## Material and Methods

### Patient tumor samples

Tumor samples were collected from patients with HNSCC undergoing surgery at François Baclesse Comprehensive Cancer Center and Caen University Hospital (France). Two regulatory frameworks were used for sample collection. Samples collected through the Biological Resource Center “BioREVA” (NF S96-900 quality management, AFNOR No. 2016:72860.5) were obtained from patients who provided written informed consent for the use of their biological material for research purposes. Additional samples were collected from patients enrolled in the ORGAVADS clinical study (IDRCB: 2019-A03332-55, NCT04261192) [22], approved by the Comité de Protection des Personnes Est III. In this framework, patients were informed that their biological samples could be used for research and were given the opportunity to express their non-opposition in accordance with French and European ethical regulations. All participants retained the right to withdraw consent or oppose the use of their samples at any time, leading to the immediate destruction of their biological material and any associated data. The histological diagnosis of HNSCC was confirmed for all samples by a certified pathologist before inclusion in the study.

### Tumor sample processing

During surgery, tumor samples were collected as part of standard medical care and transferred to the pathology laboratory. After histopathological assessment, a fragment of tumor tissue not required for diagnostic purposes was isolated and immediately placed in vials containing a collecting medium composed of Advanced DMEM (Fisher Scientific) supplemented with 10 U/mL penicillin, 10 µg/mL streptomycin, 1% GlutaMAX-1 (Fisher Scientific), 5 µg/µL Caspofungin (Merck) and 10 µM Y-27632 (Interchim). Samples were then transported at room temperature to the research laboratory. Tumor tissues were cut into approximately 4 mm^3^ fragments, which were randomly assigned to the analyses described below. One fragment was fixed in 3% paraformaldehyde for paraffin embedding and histopathological/immunohistopathological analyses. Two pieces were snapped frozen in FlashFreeze (Milestone) and stored at -150°C for subsequent DNA and RNA extraction. One fragment was used for PDTO culture, and the remaining tissue was cryopreserved in freezing solution [10% Diméthylsulfoxyde (DMSO), 90% heat-inactivated Fetal Bovine Serum (FBS)] for future isolation of viable cells. All tumor samples and derived materials were stored in the Biological Resource Center “BioREVA”.

### PDTO establishment and culture

Tumor fragment was dissociated using the human tumor dissociation kit and the gentleMACS Octo Dissociator with Heathers (Miltenyi Biotec), according to the manufacturer’s instructions. Single cells and cell clusters from tumors were collected after 100 µm filtration in organoid basal medium [OBM: Advanced DMEM (Fisher Scientific), 10 UI/mL penicillin, 10 µg/mL streptomycin, 1% GlutaMAX-1 (Fisher Scientific)] and pelleted at 430 g for 5 min. Cells were resuspended in organoid culture medium (OBM supplemented with 1X B27 (Gibco), 1.25 mM N-acetylcysteine (Sigma-Aldrich), 50 ng/mL EGF (PeproTech), 10 ng/mL FGF-10 (PeproTech), 5 ng/ mL FGF-b (PeproTech), 500 nM A-83-01 (PeproTech), 10 μM Y27632 (Interchim), 10 mM nicotinamide (Sigma-Aldrich), 1 μM PGE_2_ (PeproTech), 1 μM Forskolin (Peprotech), 0.3 μM CHIR99021 (Biogems), 100 μg/mL Primocin (InvivoGen), 50% Wnt3a/RSPO3/Noggin-conditioned medium (L-WRN, ATCC), and 10% RSPO1- conditioned medium (Cultrex HA-R-Spondin-1-Fc 293T, Amsbio). For the first 3 passages, 1 µg/mL caspofungin (Sigma-Aldrich) was added. The resulting cell suspension was mixed with 70% Cultrex Reduced Growth Factor Basement Membrane Extract, Type 2 (BME2) and seeded as 50 µL drops in a pre-warmed 24-well plate (Eppendorf). After polymerization (37°C, 5% CO2, 15 min), each drop was overlaid with 500 µL of organoid culture medium. Medium was exchanged twice a week using automated liquid handling workstation (Microlab STAR line, Hamilton). When PDTO reached around 100-150 µm in diameter, or after 4 weeks in culture, they were collected using cold OBM containing 1% Bovine Serum Albumin (OBM-BSA), centrifuged at 200 g for 2 min and incubated with TrypLE Express for up to 15 min at 37°C. Following dissociation, cells were centrifuged at 430 g for 5 min, resuspended in organoid culture medium, counted and replated at a density of 15,000 cells per 50 µl drop of 70% BME2 in pre-warmed 24-well or 6-well plates. Plates were then incubated at 37°C in a humidified 5% CO_2_ atmosphere. For biobanking, PDTO were dissociated and resuspended in Recovery Cell Culture Freezing Medium (Gibco), transferred to cryovials, and placed in a controlled-rate freezing container (CoolCell) at -80°C before long-term storage at -150°C. The identity of each PDTO was confirmed by STR profiling (Microsynth) showing concordance between the PDTO profile and that of the corresponding tumor sample.

### Histology and immunohistochemistry

Tissue samples and PDTO were fixed overnight in 3% paraformaldehyde. Fixed tissues and PDTO embedded in 2% agarose were dehydrated, paraffin-embedded, and sectioned prior to standard hematoxylin, eosin and safran (HES) staining. Immunohistochemistry was performed on paraffin-embedded tumor tissues using a Ventana Discovery Ultra (Roche) on 4-μm-thick sections. Slides were deparaffinized with UltraWash buffer at 69°C for 24 min, and epitopes were unmasked at 95°C for 56 min in CC1 buffer. Sections were incubated for 40 min at 36°C with the following primary antibodies: anti-p16 (Ventana-Roche, prediluted), anti-p53 (Abcam, 1:100), anti-p40 (Zytomed, prediluted), anti-p63 (Dako, 1:50), or anti-Ki67 antibody (Novocastra, 1:500). Detection was carried out using Omnimap Rabbit HRP secondary antibody (Ventana Medical System Inc.) for 16 min at 37°. Staining reactions were developed with 3, 3’-diaminobenzidine (DAB) and counterstained with hematoxylin using Ventana reagents, according to the manufacturer’s protocol. Slides processed without primary antibody were used as negative controls. Finally, stained slides were digitized at 20X magnification using a Vectra Polaris slide scanner (Akoya Biosciences).

### DNA and RNA extraction

DNA and RNA were extracted using the NucleoSpin Tissue® and NucleoSpin RNA kits® (Macherey–Nagel) according to the manufacturer’s instructions. Purified nucleic acids were quantified and stored at −80°C until further analysis.

### Low-Pass Whole Genome Sequencing (LP-WGS)

LP-WGS was performed on DNA samples. Library was prepared using Illumina DNA PCR-Free Prep (Illumina). Libraries were pooled all together to be sequenced on one flowcell. 150 bp paired-end sequencing of the samples was performed on the NovaSeq 6000 (Illumina). Raw reads were mapped to the human reference genome (GRCh38) using the Burrows-Wheeler Aligner (BWA) with MEM algorithm (version 0.7.17). Read duplicates were marked by Picard MarkDuplicates (version 2.27.5). Read count was performed by HMMcopy (version 0.1.1) with a bin (non-overlapping window) of 50 kb. Copy number alteration (CNA) identification and tumor fraction estimation were performed using ichorCNA (version 0.3.2). For data visualization, R/Bioconductor packages karyoploteR (version 1.16.0) and copynumber (version 1.30.0) were used.

### Targeted DNA sequencing and mutational analysis

Targeted DNA sequencing was performed by Edinburgh Genetics (Edinburgh, UK). Genomic DNA quality and quantity were assessed using the Qubit fluorometric assay (Thermo Fisher Scientific) and the QIAxcel Advanced System (Qiagen) according to the manufacturers’ instructions. Sequencing libraries were prepared using the egSEQ™ Enzymatic Library Preparation kit (Exact Genomics), followed by target enrichment with a customized egSEQ™ Pan-Cancer panel (full gene list provided in Supplementary Data File 1). Paired-end sequencing (2 × 150 bp) was carried out on an Illumina NovaSeq X Plus platform (Illumina). Raw sequencing reads were aligned to the human reference genome (hg19 build, UCSC) using BWA (version 0.7.17). Variant calling and annotation were performed using the provider’s validated bioinformatics pipeline. Variants were annotated using population frequency databases (gnomAD and the 1000 Genomes Project), gene structure annotations from UCSC hg19, and curated clinical and somatic mutation databases including ClinVar, dbSNP, COSMIC, and dbNSFP. Variants with a variant allele frequency (VAF) below 5% were excluded. Common polymorphisms with significant population frequency were filtered out based on public databases. Variants of potential functional or clinical relevance were retained for downstream analyses.

### Transcriptomic data

We processed the total RNA folowing QuantSeq 3’ mRNA-seq library kit FWD with UDI (Unique Molecular Identifiers). We used 4 µl of RNA to reach 200 ng of total RNA as input. For each sample, 1 µl of ERCC spike-in control was added. The libraries were amplified with 14 cycles. The final library is purified and checked on a High sensitivity DNA chip to be controlled on the Agilent bioanalyzer 2100. Each library is pooled equimolarly and the final pool is also controlled on Agilent bioanalyzer 2100 and sequenced on NovaSeq 6000 (Illumina) (100 bp single-end). To eliminate poor quality regions and poly(A) of the reads, we used the fastp program. We used quality score threshold of 20 and removed the reads shorter than 25 pb. The read alignments were performed using the STAR program with the human genome reference (GRCh38) and the reference gene annotations (Ensembl). The UMI (Unique Molecular Index) allowed to reduce errors and quantitative PCR bias using fastp and UMI-tools. Based on read alignments, we counted the number of molecules by gene using FeatureCount. Others programs were performed for the quality control of reads and for the workflow as qualimap, fastp, FastQC and MultiQC. Differential Gene Expression of RNA-seq was performed with R/Bioconductor package DESeq2.

### NMF on transcriptomic data

Non-negative Matrix Factorization (NMF) was applied using the butchR package. We started from a matrix of normalized expressed genes (13,401 expressed genes) and applied variance stabilizing transformation (VST). We started to apply NMF on a subset of the VST matrix subsetting to PDTO samples only. For each analysis different rank (k) were tested (from 2 to 18). Then we projected the NMF factors on all samples (tumors and PDTO) using the LCD function from the R YAPSA package using as input the count matrix (VST) and the W matrix from the NMF applied on PDTO samples. Given the NMF on all samples, we plotted UMAP projection using ggplot and computed distance between paired samples (PDTO/tumor) based on the UMAP. To choose the best rank, we choose the one that minimize the sum of distances between paired samples (PDTO/tumor). We then fixed the rank to k=18. Each NMF-derived signature corresponds to a latent transcriptional program characterized by a specific gene weight distribution.

### PDTO treatment

When PDTO reached a diameter of 75-150 µm, they were collected with cold OBM-BSA and centrifuged at 200 g for 2 min. Pelleted PDTO were resuspended in organoid treatment medium (organoid culture medium lacking primocin, Y-27632 and N-acetylcysteine) and counted. Subsequently, they were diluted in 2% BME2/organoid treatment medium, and 200 PDTO per well were seeded in 100 μL in white, clear-bottom 96-well plates (Greiner) previously coated with a 1:1 mixture of organoid treatment medium and BME2. For radiotherapy response assays, PDTO were irradiated in microtubes using the CellRad irradiator (FAXITRON Bioptics) prior to plating, and medium was replaced after 5 days. For chemotherapy response assays, drugs were prepared in 2% BME2/organoid treatment medium and added 1 hour after PDTO plating. PDTO morphology was monitored using IncuCyte S3 (Sartorius). After 7 days (chemotherapy) or 10 days (radiotherapy), ATP levels were quantified using CellTiter-Glo 3D assay (Promega), and luminescence was measured with the GloMax reader (Promega). Cell viability values were normalized to control. Two independent biological replicates were realized in most cases and treatment sensitivity was determined based on the earliest available passage of each PDTO model. A positive control of citotoxicity (cycloheximide 75 µM) was included used in cisplatin assays to calculate a z-score as follows:

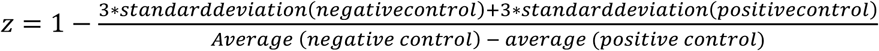

Experiments with a z-score below 0.4 were considered non-exploitable. Each experiment was performed at least twice (n=2). Viability curves were generated using GraphPad Prism (v9.2.0), the area under the curve (AUC) was computed for each PDTO model. Normalized AUC z-scores were calculated by subtracting the mean AUC from each value and dividing by the standard deviation across all models.

### Statistical analysis

Categorical variables were described using frequencies and percentages, whereas continuous variables were displayed as mean ±SD or median and range. Patient characteristics were compared between samples that resulted in the establishment of a long-term PDTO and those that did not, using the chi-square test (or Fisher exact test when expected frequencies were <5) for categorical variables, and the Student’s t-test (or Wilcoxon-Mann-Whiney test for non-gaussian distributions) for quantitative variables. All statistical tests were two-tailed, and a p value <0.05 was considered statistically significant, while a p value <0.10 was considered a relevant trend. The association between long-term PDTO establishment and patient characteristics was assessed using univariate logistic regression models. The association between PDTO viability under different treatments and the clinical response of patients at 24 months was assessed using logistic regression models. Receiver operating characteristic (ROC) analysis was conducted to determine the optimal cut-off value for viability measurements in predicting patient responses at 24 months, based on the Youden index. Survival rates were estimated using the Kaplan Meier method and survival distributions were compared using the log-rank test.

## Results

### Establishment of a panel of PDTO from head and neck cancer

HNSCC tumors samples were collected during the initial surgery and then processed according to our previously published protocol [22], as described in Materials and Methods section. Both tumors and established PDTO lines were biobanked as frozen and paraffin embedded specimens for subsequent histological and molecular characterizations, and cryopreserved as viable cells. PDTO were considered successfully established long-term lines when genetic filiation with the parental tumor was validated by STR profiling, malignant features were confirmed and sustained proliferation was maintained beyond 7 passages. PDTO which failed to maintain proliferation beyond 7 passages were called “short-term PDTO”. Established PDTO lines were subsequently exposed to standard-of-care treatments for HNSCC to evaluate treatment sensitivity, which was compared with the clinical response of the corresponding patients to assess their predictive potential (**Figure 1A**). Samples were collected from 109 patients. Two cases were diagnosed as either *in situ* carcinoma or adenocarcinoma on definitive anatomopathological analysis and were excluded *a posteriori* from the study (**Figure 1B**). Among the remaining 107 patients, patient-derived organoids (PDO) formation at passage 0 (PDO p0) was observed in 77 samples (72%). Six of these cultures were lost due to contamination or technical issue and 22 failed to proliferate beyond the initial passages. Forty-nine sample generated PDO beyond passage 3; among them 15 of them were identified as non-malignant organoids either by histological analysis or genetic analysis and were excluded from further analysis (**Figure S1**), while 14 failed to generate PDTO beyond passages 4 to 6. Overall, 20 long-term PDTO lines were successfully established, corresponding to a global establishment rate of 18.7% (**Figure 1C**). The rate of establishment during the first year of the project was 8,79%. Over the next three years, the establishment rate of long-term PDTO increased to 21,4% following optimization of culture conditions, including media composition and extracellular matrix modifications (the optimal protocol is detailed in Materials and Methods) (**Figure S2**).

**Figure 1.**
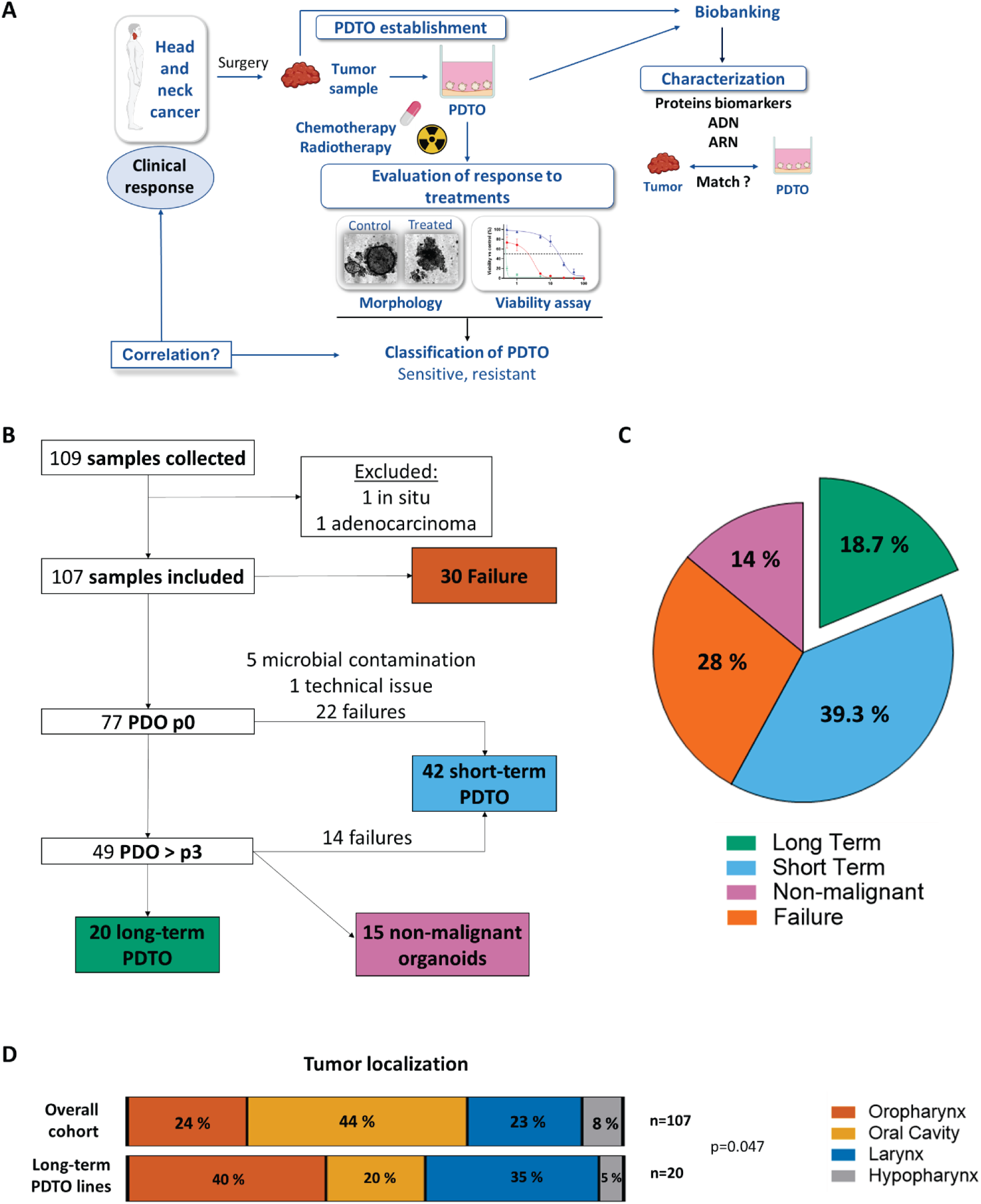
Establishment of a panel of PDTO from HNSCC. **(A)** Overview of the study design. **(B)** Flowchart showing successful and unsuccessful establishment of PDTO cultures. Long-term PDTO defined as : genetic filiation with the parental tumor validated by STR profiling, malignant features confirmed, sustained proliferation maintained beyond 7 passages. Short-term PDTO defined as : malignant features confirmed, sustained proliferation maintained beneath 7 passages. **(C)** Establishment rate among the 107 patients included in the overall cohort. **(D)** Anatomical distribution of tumors in the overall patient cohort compared with tumors giving rise to long-term PDTO lines.

Long -term PDTO were derived from patients with a mean age of 62 years, of whom 90% were male. Three PDTO lines originated from HPV-positive tumors (oropharyngeal cancers, p16-positive). Four patients had a history of previous HNSCC treated with radiotherapy alone or in combination with surgery and/or chemotherapy. Clinical characteristics of the 20 patients from whom long-term PDTO were established PDTO are summarized in **Table S1**.

The anatomical distribution of tumor sites was similar between the overall cohort of patients and the long-term PDTO cohort, with the exception of a lower representation of oral cavity tumors among successfully established PDTO (**Figure 1D**). No statistically significant differences were observed between successful and unsuccessful establishment groups with respect to age, sex, previous HNSCC, or post-operative treatment (**Table S2**). These parameters, as well as tumor differentiation grade, tobacco and alcohol consumption and cancer cell content in the sample, were not associated with successful establishment of either short-term or long-term PDTO (**Table S3**).

### PDTO recapitulate histology and tumor marker expression

HNSCC-derived PDTO displayed a relatively homogenous morphology. They exhibited spherical architecture, high cellular density, and a refractile outer border (**Figure 2A**). No cystic morphology was observed. Histological features and HNSCC-specific biomarkers were compared between tumor and their corresponding PDTO lines. Representative HES staining and immunohistochemical analyses from two models, one HPV-positive and one HPV-negative, are shown in **Figures 2B–C**, with additional models presented in **Figure S3**. Key histological characteristics of HNSCC, including a high nuclear--to--cytoplasmic ratio, nuclear pleomorphism, and prominent nucleoli, were preserved in PDTO. Well-defined tumor cell borders, intercellular bridges and keratinization were also observed in some cases, as shown in **Figure 2B**. HNSCC-associated biomarkers such as p40 and p63 were detected in both tumors and matched PDTO lines, with nuclear immunostaining observed in all specimens, as expected (**Figure 2C**). p53 expression was variable but showed concordant staining patterns between tumor and paired PDTO. p16, a surrogate marker of HPV-positive HNSCC, was strongly expressed in the 3 oropharyngeal tumors and paired PDTO compared to HPV-negative samples.

**Figure 2.**
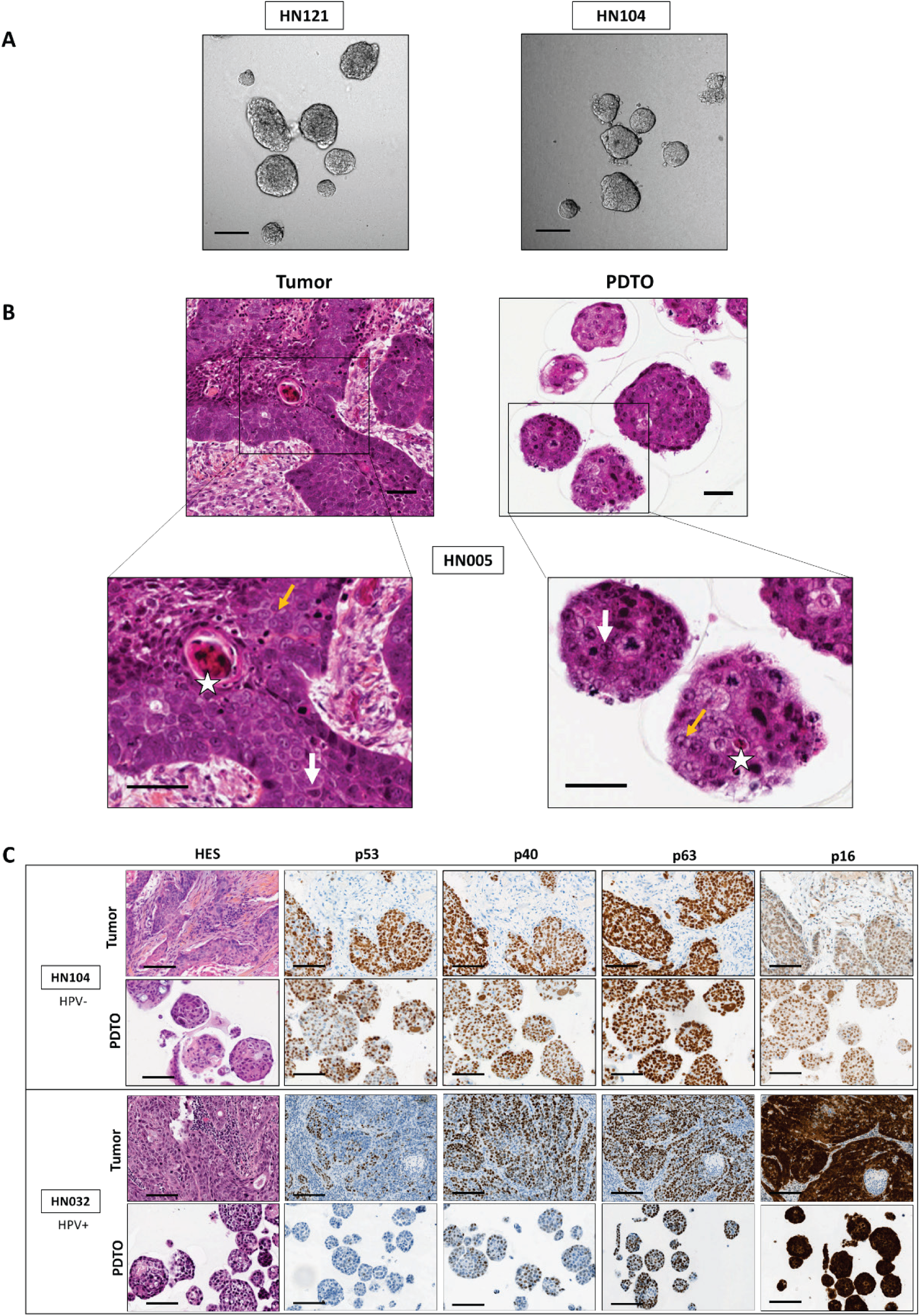
Histology and tumor marker expression in PDTO and paired tumors. **(A)** Representative brightfield images of PDTO. Scale bar: 2Q0 pm. **(B)** Hematoxylin-eosin-saffron (HES) staining of the PDTO line HN005 and its paired tumor, showing characteristic histological features of HNSCC. Star: keratin pearl; white arrow: well-defined cell borders; yellow arrow: irregular nuclei with prominent nucleoli. Scale bar: 50 pm. **(C)** Immunohistochemical analysis of HNSCC-associated biomarkers (p!6, p53, p40, and p63} in two PDTO lines and their paired tumors (one HPV-positive and one HPV-negative). Scale bar: 100 pm.

### Genetic characteristics of the tumor of origin are preserved in PDTO over time

Copy-number variations (CNVs) across the genome of 16 PDTO and their matched primary tumors were analyzed using low-pass whole-genome sequencing (LP-WGS) (**Figure 3A**). Overall, PDTO CNV profiles closely mirrored those of their tumors of origin, with an average similarity score of 0.65 (**Figure 3B**). Cases displaying lower similarity (e.g., HN-099) were associated with the absence of detectable alterations in the primary tumor sample, likely reflecting a high proportion of normal stromal and immune cells within the tumor specimen (e.g. only 15% of tumor cells in the primary tumor sample of HN-099). Analysis of the most frequently mutated genes in HNSCC [23] revealed a strong concordance between PDTO and their corresponding tumors (**Figure 3C**). Importantly, these genetic and genomic alterations were stably maintained across serial passages and following cryopreservation, as indicated by Dice–Sørensen coefficients ranging from 0.8 to 1 (**Figure S4A–C**). Furthermore, CNV profiling enabled the reclassification of three PDTO models as non-malignant organoids, based on the absence of detectable genomic alterations (**Figure S5A**). This finding was subsequently validated by immunohistochemistry, which revealed the presence of proliferative basal cells (Ki67- and TP63-positive) at the periphery of these organoids (**Figure S5B**). Together, these results demonstrate that PDTO faithfully preserve the genomic landscape of their tumors of origin over time.

**Figure 3.**
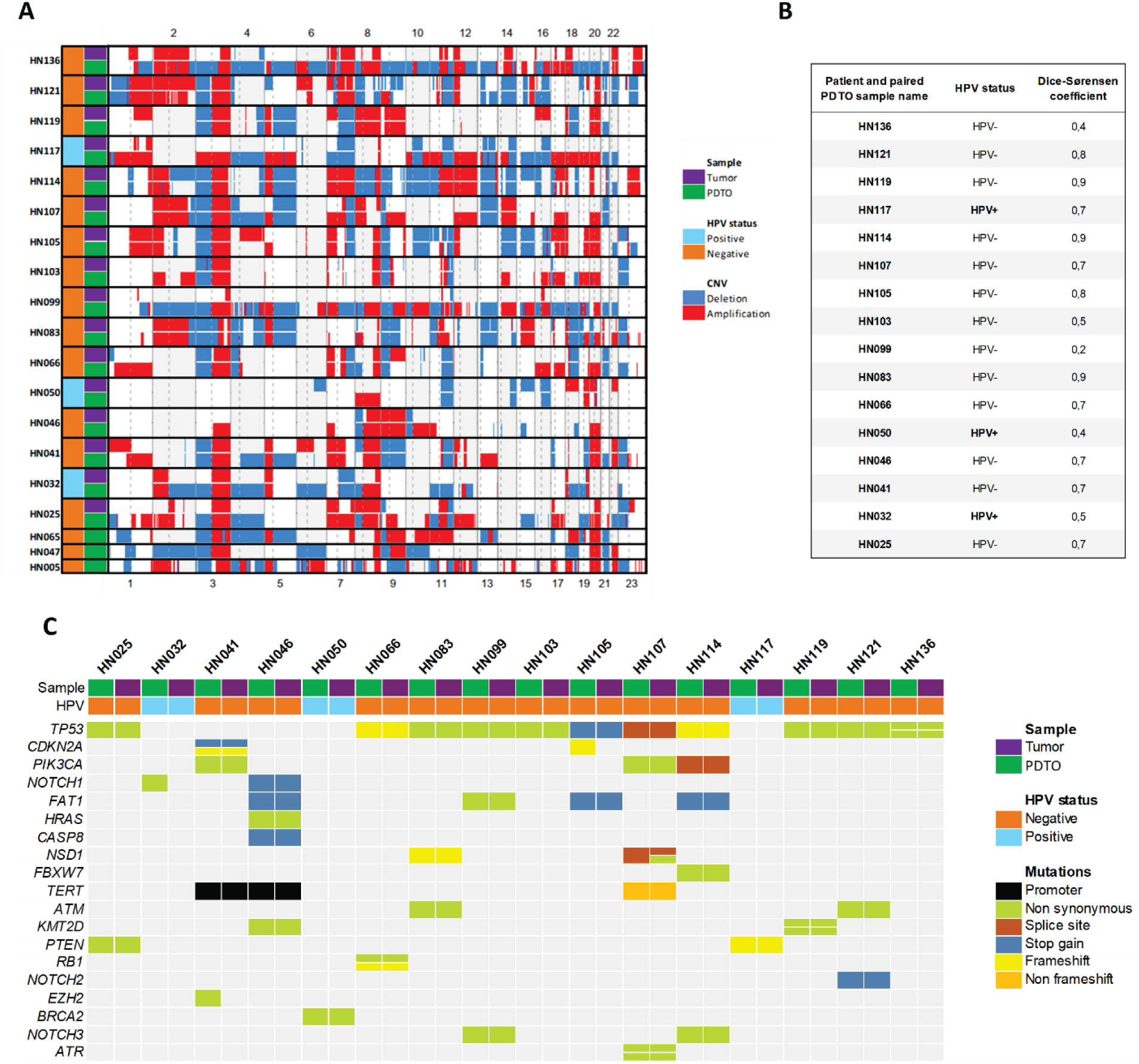
Genetic characteristics of PDTO and their tumors of origin. **(A)** Copy-number variation (CNV) profiles analyzed by low-pass whole-genome sequencing (LP-WGS) in 16 PDTO and their paired tumors. **(B)** Correlation of CNV profiles between PDTO and paired tumors, calculated using the Dice-S0rensen coefficient **(C)** Mutation status of 19 major genes frequently mutated in HNSCCin 16 PDTO and paired tumors.

#### Transcriptomic profiling confirms the overall concordance between PDTO and their tumors of origin

Transcriptomic analyses were performed on paired tumor samples and PDTO (n = 15). Gene expression levels showed a strong correlation between each PDTO and its corresponding tumor, with R values ranging from 0.90 to 0.96 (**Figure 4A**). Despite this overall concordance, Uniform Manifold Approximation and Projection (UMAP) of all samples revealed a clear separation between primary tumors and PDTO, with tumor samples clustering together and distinct from PDTO profiles (**Figure 4B**). As expected, functional enrichment analysis of differentially expressed genes indicated a strong overrepresentation of immune-related pathways in tumor samples, consistent with the presence of stromal and immune components in the original tissue and their absence in PDTO cultures (data not shown). To enable a more refined comparison of matched samples and to focus on dominant transcriptional signals, we applied a non-negative matrix factorization (NMF)-based approach, as previously described in our recent work [24]. This method decomposes transcriptomic profiles into a limited set of latent transcriptional programs, allowing each sample to be represented by its relative contribution to these programs, thereby reducing technical and biological noise. Using these NMF-derived representations, UMAP analysis revealed a strong concordance between PDTO and their tumors of origin (**Figure 4C**), as highlighted by the marked decrease in distance between paired samples (**Figure 4D**). Consistently, unsupervised clustering showed a tendency for tumor–PDTO pairs to cluster together (**Figure 4E**). Transcriptomic profiles were also conserved across passages, as illustrated for models HN-005, HN-041, and HN-114 (**Figure 4C**). Collectively, these analyses demonstrate that, despite the absence of microenvironmental components, PDTO retain the major transcriptional features of their tumors of origin.

**Figure 4.**
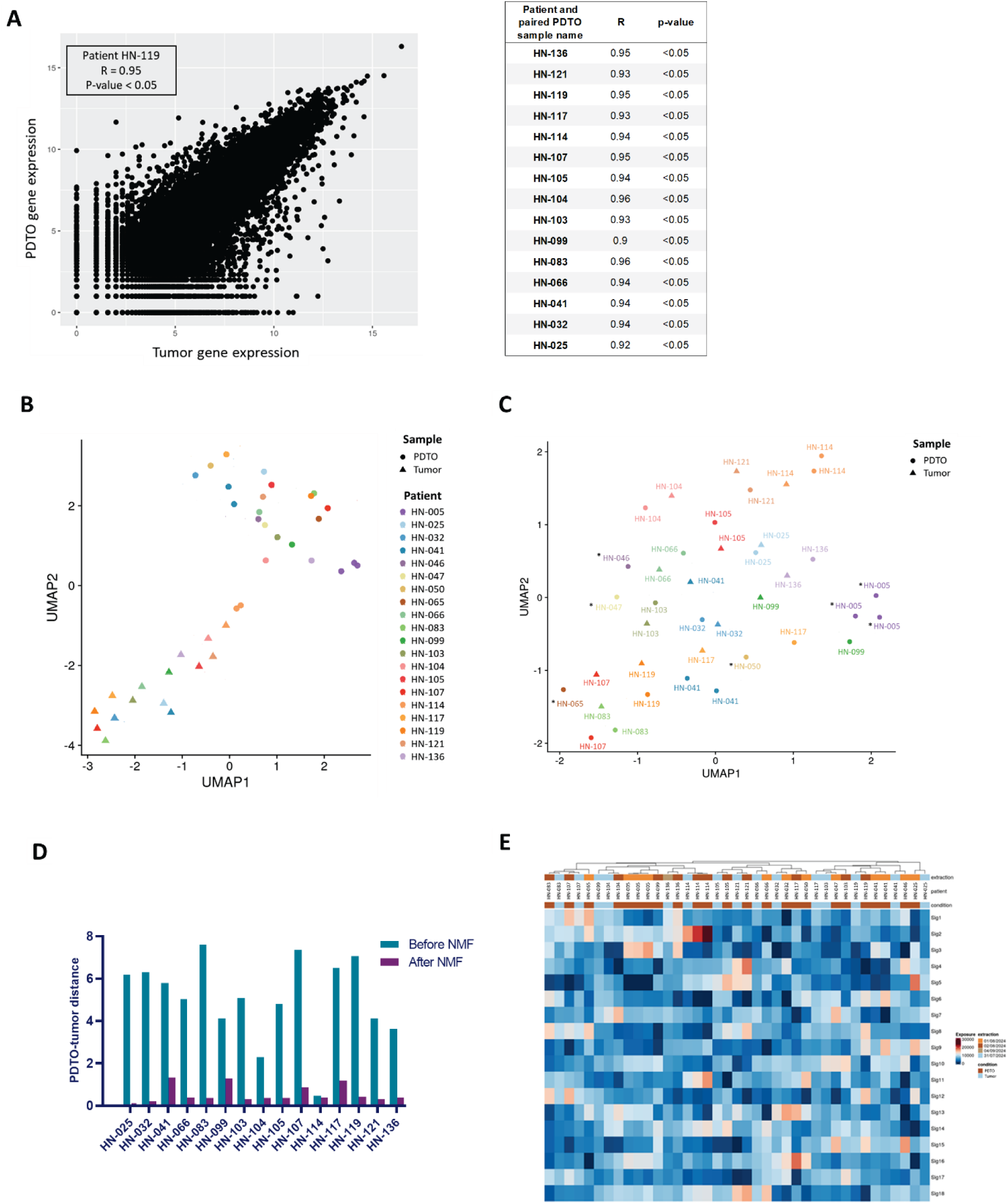
PDTO recapitulate the cancer cell gene expression programs of their tumors of origin. **(A)** Scatter plot showing gene expression levels in a representative tumor-PDTO pair (patient HN-119, left), and Pearson correlation coefficients between each tumor and its matched PDTO across the cohort (right). **(B-C)** Distribution of tumor samples and paired PDTO across Uniform Manifold Approximation and Projection (UMAP) clusters based on gene expression profiles **(B)** before and **(C)** after non-negative matrix factorization (NMF, k = 18). *PDTO without a matched primary tumor sample. **(D)** Histogram showing the distribution of Euclidean distances between tumor samples and their paired PDTO using UMAP coordinates, before and after NMF (k = 18). **(E)** Hierarchical clustering and heatmap of exposure values for PDTO and tumor samples, based on non-negative matrix factorization (k = 18) of RNA-seq expression data.

### PDTO showed heterogeneous response to treatments

PDTO lines were exposed to increasing concentrations or doses of standard-of-care post-operative treatments for HNSCC (cisplatin and X-rays), and cell viability was assessed using biochemical assay. Marked heterogeneity in treatment response was observed across PDTO lines for both cisplatin and irradiation. Cisplatin IC50 varied by approximately two orders of magnitude between the most sensitive and the most resistant models. Similarly, a tenfold difference in sensitivity was observed following X-ray exposure.

In response to cisplatin, five PDTO lines exhibited marked sensitivity compared with the others, with viability decreasing to nearly 0% at 5 µM (**Figure 5A**). Beyond these highly sensitive models, dose-response curves were continuous, making it difficult to define discrete subgroups. Notably, HPV+tumor-derived models were among the most sensitive, consistent with the known clinical responsiveness of these tumors. Morphological observations were consistent with viability results. In control conditions, PDTO increased in size over time, whereas exposure to 10 µM cisplatin inhibited growth in a sensitive model (HN-121) (**Figure 5B**). Under cisplatin treatment, sensitive PDTO (HN-121) appeared disaggregated. In contrast, resistant PDTO (HN-099) maintained their size or continued to grow at 10 µM.

**Figure 5.**
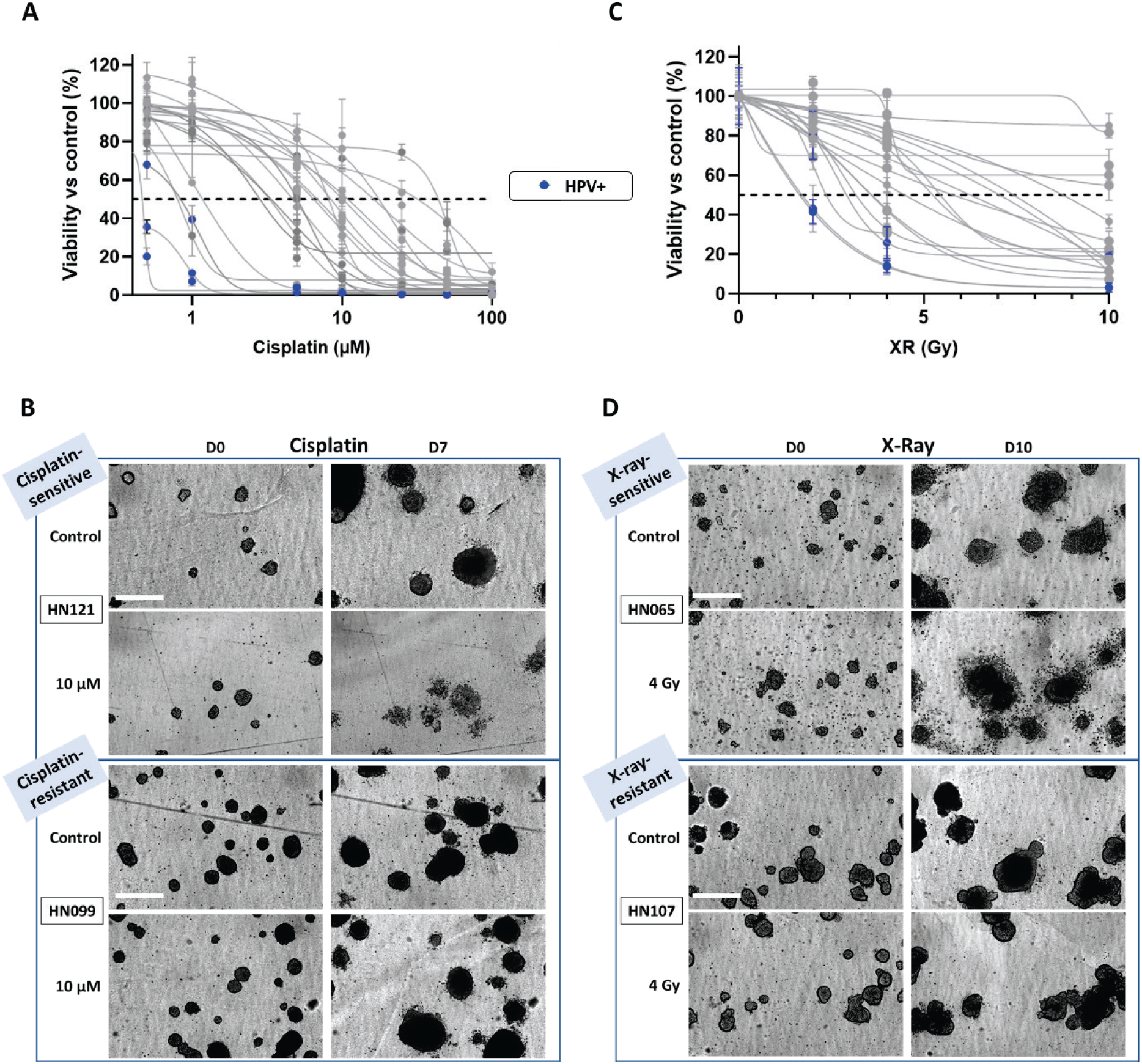
Response of PDTO lines to conventional treatments. (n = 20) Dose-response curves of viability (logarithmic scale) of PDTO lines following exposure to increasing **(A)** concentrations of cisplatin or **(C)** doses of X-rays. Representative morphology of a sensitive and a resistant PDTO line following **(B)** cisplatin or **(D)** X-ray exposure. Blue curves correspond to HPV-positive PDTO. Scale bar: 400 pm.

Following X-ray exposure, five PDTO lines exhibited high resistance, with viability remaining above 50% at 10 Gy (**Figure 5C**). At 4 Gy, two groups could be distinguished: one with viability above 50%, including four of the previously identified resistant lines, and another with viability below 50%. Again, HPV+ tumor-derived models were among the most sensitive. Morphological changes were consistent with these findings. The sensitive PDTO line (HN-065) showed structural disruption after exposure to 4 Gy, although some PDTO continued to grow at this dose. In contrast, resistant PDTO (HN-107) retained a morphology similar control condition, with continued growth and well-defined borders (**Figure 5D**).

### PDTO responses predict clinical outcome

Area under the curve (AUC) values derived from dose–response analyses were used to classify PDTO according to their sensitivity to cisplatin and X-rays (**Figure S6A–B**). Clinical status was annotated at 9, 18, and 24 months. Although isolated discrepancies were observed, heatmap visualization revealed an overall concordance between PDTO drug responses and patient outcomes. For cisplatin, PDTO response was significantly associated with clinical status at 24 months (p = 0.012) and with disease-free survival (DFS) (p = 0.0054). Similar trends were observed for X-rays, with associations approaching statistical significance for 24-month response (p = 0.092) and reaching significance for DFS (p = 0.05). Restricting the analysis to patients who received post-operative radiotherapy attenuated statistical significance, likely due to reduced sample size (n=15). PDTO responses to cisplatin and X-rays were correlated (Pearson r = 0.6215, 95% CI: 0.2469–0.8345; p = 0.034) (**Figure S6C**), showing partially overlapping sensitivity profiles. Receiver operating characteristic (ROC) analyses were performed to determine optimal thresholds (**Figure 6A–C**). The area under the ROC curve was 0.8021 for cisplatin and 0.7604 for X-rays, indicating acceptable discriminatory performance. Using the optimal Youden index cut-off, cisplatin sensitivity was predicted with 66.7% sensitivity [95% CI: 39.06-86.19%] and 100% specificity [95% CI: 67.56-100%], while X-ray sensitivity reached 91.7% [95% CI: 64.61-99.57%] with 75% specificity [95% CI: 40.93-95.56%]. Survival analyses confirmed that PDTO response was significantly associated with DFS for both cisplatin (p = 0.0022) and X-rays (p = 0.0020) (**Figure 6B–D**). Notably, this association remained significant in patients who received post-operative radiotherapy (p = 0.015), despite the limited cohort size (n = 15) (**Figure S6D**).

**Figure 6.**
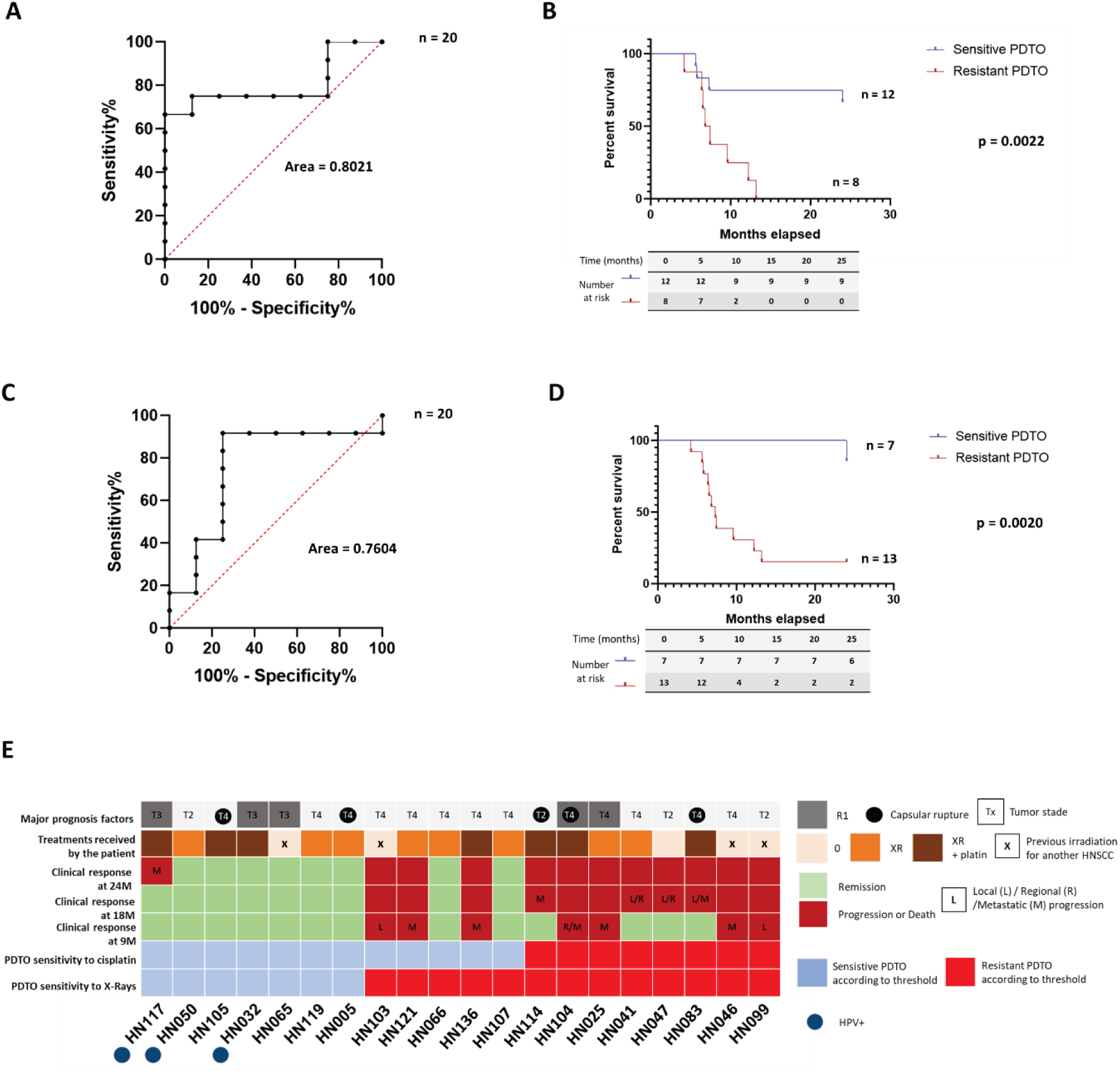
Association between PDTO treatment sensitivity and clinical outcome. ROC curve constructed using PDTO response to **(A)** cisplatin and **(C)** X-rays (AUC) and clinical response at 24 months (good vs. poor responders). The optimal threshold was determined using the Youden index. Kaplan-Meier curves of disease-free survival (DFS) stratified according to PDTO sensitivity to **(B)** cisplatin and **(D)** X-ray based on the ROC-defined threshold. **(E)** Summary heatmap integrating clinical response at different time-points, postoperative treatment received, major histopathological prognostic factors and sensitivity classification (according to AUC values for X-ray and cisplatin response).

Stratified analyses further highlighted the clinical relevance of dual-modality sensitivity. All patients whose PDTO were classified as resistant to both cisplatin and X-rays relapsed within 18 months (**Figure 6E**). In contrast, patients whose PDTO were sensitive to both modalities maintained durable disease control, with only one relapse observed at 24 months (**Figure 6E**).. Patients with discordant sensitivity profiles exhibited intermediate and more heterogeneous outcomes. Collectively, these findings demonstrate that *in vitro* PDTO responses to cisplatin and radiotherapy capture clinically meaningful differences in patient prognosis, supporting their potential as predictive functional biomarkers in HNSCC.

## Discussion

We successfully established 20 PDTO lines from 107 patients, maintained over 7 passages and cryopreserved. Our global cohort of 107 patients is representative of the usually described population of HNSCC with a majority of men, a mean age between 60-65 years, and a classical localization distribution with a majority of oral cavity and oropharynx and a smaller amount of larynx and hypopharynx [25]. Our cohort of 20 patients with long-term PDTO was representative of this global cohort.

The long-term establishment rate following culture optimization was 21.4%. Our results are in agreement with the ones described by Millen *et al.* (28.4%) [16]. Other studies described establishment rates for PDTO derived from HNSCC from 30% [17] to 92,8% [15]. However, the definition of established PDTO line is variable and generally based on 3 passages or even on the obtention of PDTO at passage 0 (p0), which would correspond to establishment rate of 45.79% or 71.96%, respectively, in our cohort. Unlike those studies, we also chose to include all our samples in the denominator including the samples that led to non-malignant organoid generation, the samples with a small amount of tumor cells and the samples with microbial contaminations. Taking all these samples into account allows for a more accurate assessment of the actual proportion of patients for whom predictive functional assays could have been performed in a clinical setting.

No prognostic factor of establishment of long-term PDTO or of PDO p0 appearance was found in our study. Some have suggested a better establishment for younger patients [13] and more precisely under 68 years old [26] but we have not confirmed that in our cohort. The amount of epithelial cells in the initial sample or after dissociation has been described to improve establishment [16, 26]. No correlation was found in our study with the proportion of cancer cells on histological analysis (data not shown), although the latter does not directly quantify epithelial cell content.

The optimization of culture method allowed an improvement of the establishment rate for long term PDTO from less than 9% to more than 20%. This evolution coincided with the emergence of non-malignant organoids: either non-malignant organoids were more effectively distinguished from short-term organoids in the second period, or the culture conditions that promoted the growth of long-term PDTO also promoted the expansion of non-malignant organoids. The culture conditions commonly used for HNSCC organoids may indeed support the growth of normal epithelial organoids [12], which represents a potential limitation in the context of functional precision medicine. A higher success rate of establishment of non-malignant organoids over PDTO has already been reported [27]. Given that oral cavity is the primary HNSCC site, minimizing contamination by non-malignant organoids is critical. In our cohort, non-malignant organoids were identified in 15 cases, highlighting the necessity of early detection and exclusion. Currently, their identification relies on histological aspect of cells, immunohistochemical features, namely proliferative basal cells (Ki67- and TP63-positive) located at the periphery [12, 27], or on the absence of tumor-specific genomic alterations, such as copy-number variations. However, both approaches require additional expansion of biological material and processing time, which may delay downstream functional assays and limit compatibility with rapid, clinically actionable precision-medicine workflows. Pharmacological selection using Nutlin-3 has been proposed to enrich for *TP53*-mutated cancer cells [28]. However, as *TP53* mutations are not universal in HNSCC, this strategy may introduce a selection bias toward specific tumor subclones. An alternative approach would involve careful macroscopic selection of tumor tissue by the surgeon or pathologist to minimize contamination with normal epithelium. Currently, non-malignant and tumor-derived organoids may display similar morphological features, especially in the oral cavity [29], making morphology-based discrimination unreliable. Nevertheless, in the future, automated image-based classification using artificial intelligence, potentially coupled with organoid-picking platforms, may offer a promising strategy to identify and eliminate non-malignant organoids in a standardized and scalable manner.

The PDTO lines histological analysis recapitulates the tumoral features of HNSCC and the immunohistochemical markers of the original tumor are conserved. We established three HPV+ PDTO lines, derived from p16 positive oropharyngeal cancer. Others described until 11 HPV+ PDTO lines [13]. HPV+ models cell lines available are not numerous and having new models of this subtype of HNSCC is important to pursue preclinical research [30].

Genetic analyses demonstrated that PDTO faithfully recapitulate both the CNV landscape and the mutational profile of the original tumors, and that these genomic features are maintained across serial passages and after cryopreservation. As expected, the major alterations commonly described in HNSCC were identified in most of our tumor samples and corresponding PDTO, including amplifications of 3q, 5p, and 8q, as well as deletions of 3p and 5q [31]. Similarly, *TP53* mutations were detected in 11 of 13 (84%) HPV-negative PDTO/tumor pairs, a frequency similar to that reported by TCGA (86%) [32], whereas no *TP53* mutation was identified in the three HPV-positive pairs (TCGA: 0.02%). Those results are consistent with our immunohistochemistry analysis where 4 of the 5 PDTO lines without *TP53* mutation presented a wild-type pattern with some scattered positive cells where mutated PDTO lines presented mostly pathological pattern with stronger staining or absence of staining [33]. In addition, transcriptomic analysis based on a non-negative matrix factorization (NMF) approach enabled us to directly compare the cancer cell component of primary tumors with that of the corresponding PDTO, without requiring prior microdissection nor single cell approaches. Consistent with our previous observations in ovarian PDTO [24], the transcriptomic profile of cancer cells from the original tumor is faithfully preserved in the derived PDTO. Altogether, these results highlight the strong biological relevance of our PDTO models, which faithfully preserve the genomic and transcriptomic landscape of the original tumors. These findings further support their value as robust preclinical models and underscore the importance of prospectively evaluating their predictive potential in the context of precision oncology.

PDTO lines displayed heterogeneous responses to cisplatin and X-rays, as expected and as reported in previous studies [11, 16, 34], supporting their ability to capture the clinical heterogeneity of treatment response in HNSCC. Importantly, PDTO-based stratification revealed prognostic information that was not reflected by classical clinicopathological parameters such as nodal extracapsular extension, R1 resection status, or tumor volume, which were evenly distributed among good and poor PDTO responders. Notably, cisplatin sensitivity of PDTO was strongly associated with disease-free survival (DFS) across the entire cohort, regardless of whether patients received adjuvant radiotherapy or cisplatin-based radiochemotherapy. These findings are consistent with recent evidence demonstrating that cisplatin responsiveness of HNSCC PDTO correlates with clinical outcomes in patients treated with adjuvant chemoradiotherapy [20]. We also observed a trend toward correlation between PDTO radiosensitivity and long-term clinical outcome (follow-up >24 months). Although limited by cohort size, our results align with several independent studies. Driehuis *et al.* reported that higher AUC values of viability curves were associated with increased relapse risk in a cohort of seven patients [12]. Similarly, radiotherapy response of HNSCC PDTO has been linked to clinical outcome in cohorts of 8 to 15 patients [16, 19], with GR50 values correlating with adjuvant radiotherapy response. Lee *et al.* also observed improved progression-free survival in patients whose PDTO displayed lower AUC in response to irradiation [8], although only two-thirds of their cohort received postoperative radiotherapy. Finally, the biological relevance of our model is further supported by the behavior of HPV-positive PDTO, which were among the most treatment-sensitive lines for both cisplatin and X-rays. This observation is consistent with the well-established favorable prognosis of HPV-positive HNSCC, likely related to increased sensitivity to chemoradiotherapy [6].

All patients whose PDTO were classified as resistant to cisplatin and X-rays experienced relapse within 18 months, whereas all patients whose PDTO were classified as sensitive to both modalities showed durable clinical response at 18 months. However, these findings should be interpreted cautiously, as they reflect a prognostic association rather than definitive predictive evidence. Indeed, not all patients received postoperative radiochemotherapy, and surgical management may have influenced outcomes. Validation in a prospective cohort of patients treated with primary radiotherapy or radiochemotherapy will therefore be essential to eliminate the confounding effect of surgery and formally assess predictive value. Nonetheless, the fact that all patients with PDTO resistant to both modalities relapsed suggests potential clinical utility, potentially guiding early consideration of innovative therapies or trial enrollment.

Despite the strong overall concordance between *in vitro* PDTO sensitivity and clinical response, a few discrepancies were observed. For instance, the HN-107 PDTO displayed radioresistance, while the corresponding patient achieved sustained disease control with no relapse after 24 months. Notably, this PDTO line remained sensitive to cisplatin. Importantly, this patient underwent complete surgical resection, which may have significantly contributed to the favorable outcome and thus confounded the interpretation of radiotherapy response. Another potential explanation for such discrepancies relates to one of the main limitations attributed to PDTO models, the absence of a tumor microenvironment, which is known to influence treatment response [35]. Recent studies have developed co-culture systems combining HNSCC PDTO with cancer-associated fibroblasts (CAFs) [18, 36], demonstrating reciprocal phenotypic modulation of both fibroblasts and tumor cells. Whether incorporation of CAFs or other stromal and immune components could further enhance the predictive performance of PDTO remains to be determined. Nevertheless, the strong association observed in our cohort between PDTO sensitivity and clinical outcome suggests that, at least in this setting, a cancer-cell–intrinsic model may already capture a substantial proportion of clinically relevant therapeutic vulnerability. Increasing model complexity might further improve predictive power but could also reduce overall applicability and throughput.

Correlation between PDTO drug response and clinical outcome has already been reported in other tumor types, particularly in colorectal cancer [10]. The growing interest in using PDTO as predictive “avatars” to guide individual patient treatment has led to several prospective clinical trials [37, 38], which have also highlighted important practical and methodological challenges. One major limitation is the establishment rate. In our study, long-term PDTO lines were successfully generated in approximately 20% of cases. However, organoids at p0 were obtained in nearly 70% of cases. This suggests that if treatment sensitivity assays could be miniaturized and optimized to operate on small amounts of material at early passages, PDTO-based testing could become feasible for a substantially larger proportion of patients. Similar high establishment rates have been reported using miniaturized pillar/well systems, reaching over 90% efficiency [15]. A key prerequisite for such early-stage applications will be the rapid discrimination between non-malignant and tumor-derived organoids, as discussed above. Miniaturization would also reduce the turnaround time required to assess treatment sensitivity, thereby improving compatibility with clinical decision-making timelines. Moving beyond the postoperative setting and integrating PDTO-based testing earlier in the treatment pathway would likely require the use of diagnostic biopsies. Encouragingly, recent work has demonstrated that PDTO can be established from HNSCC biopsy specimens without reduced efficiency compared to surgical samples [13].

Beyond their potential for predicting clinical response, HNSCC-derived PDTO have already been employed in preclinical studies to evaluate innovative therapeutic strategies [39–43]. However, large-scale and high-throughput implementation of PDTO technology will be essential to fully exploit PDTO biobanks for drug screening and therapeutic discovery. In this context, imaging-based response assessment, rather than traditional viability assays, may represent a promising strategy to enhance scalability and analytical efficiency [41, 44]. The development of such high-throughput platforms will be critical to streamline PDTO integration into precision oncology pipelines.

## Conclusion

In conclusion, our work strengthens the rationale for using HNSCC-derived PDTO as biologically faithful and clinically relevant models for both preclinical research and precision oncology. We demonstrate, in the largest clinically annotated HNSCC PDTO cohort to date, a robust association between *ex vivo* sensitivity to cisplatin and X-rays and patient prognosis. These findings position PDTO not only as experimental surrogates of tumor biology, but as promising functional tools to refine therapeutic stratification and improve treatment selection. The continued development of standardized, miniaturized, rapid, and reproducible workflows will be essential to ensure compatibility with clinical timelines. Equally important will be the identification of clinical contexts in which PDTO-based functional assays provide maximal added value, thereby enabling their optimal integration into the management of patients with HNSCC. In parallel, PDTO represent powerful tools for the research community to investigate disease biology, evaluate novel therapeutic strategies, and explore treatment response and resistance at single-cell resolution.

## Acknowledgments

The authors thank Adèle Riot (Department of Biopathology, Comprehensive Cancer Center François Baclesse) for her assistance with histological analyses, and Jérôme Toutain (Cyceron core facility, University of Caen) for X-ray experiments. We also thank Benoît Goudergues Biological Resource Centre “BioREVA”, Comprehensive Cancer Center François Baclesse), as well as Edwige Abeilard and Solène Angot (INSERM U1086 ANTICIPE, Université de Caen Normandie) for their valuable technical support. We are grateful to Emmanuelle Brouart, Valerie Marie-Couzinet and Pascale Marie Zeimet (Calvados General Tumor Registry, Comprehensive Cancer Center François Baclesse) for clinical data collection. We also thank Nathalie Rousseau (Biological Resource Centre “BioREVA”, Comprehensive Cancer Center François Baclesse) for her support with biological sample storage, and Christophe Denoyelle (ImpedanCELL core facility, Service Unit PLATON, University of Caen Normandie) for real-time imaging experiments. The IncuCyte S3 device was acquired with the support of the French State and the Normandy Regional Council (Contrat de Plan État–Région, CPER INNOVONS). The ORGAPRED project is co-funded by the Normandy Regional Council and the European Union within the framework of the ERDF/ESF 2014–2020 Operational Programme, conducted as part of the 2015–2020 State–Region Planning Contract.

## Availability of data and materials

NGS, CNA and RNA-seq data will be made available to the community (on GEO or equivalent) upon publication. During the review process, these data will be made available to the reviewers and editor upon request. All other data supporting the findings of this study are available from the corresponding authors on reasonable request. All materials will be available upon request through a material transfer agreement.

---

**Figure S1.**
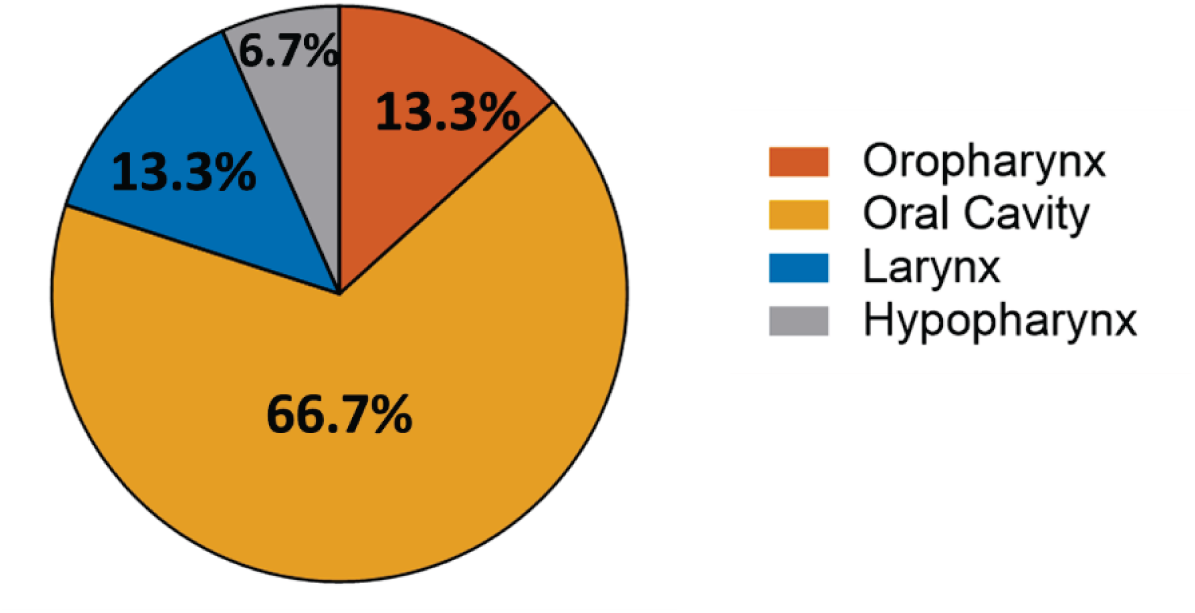
Anatomical origin of samples giving rise to normal organoids (n = 15).

**Figure S2.**
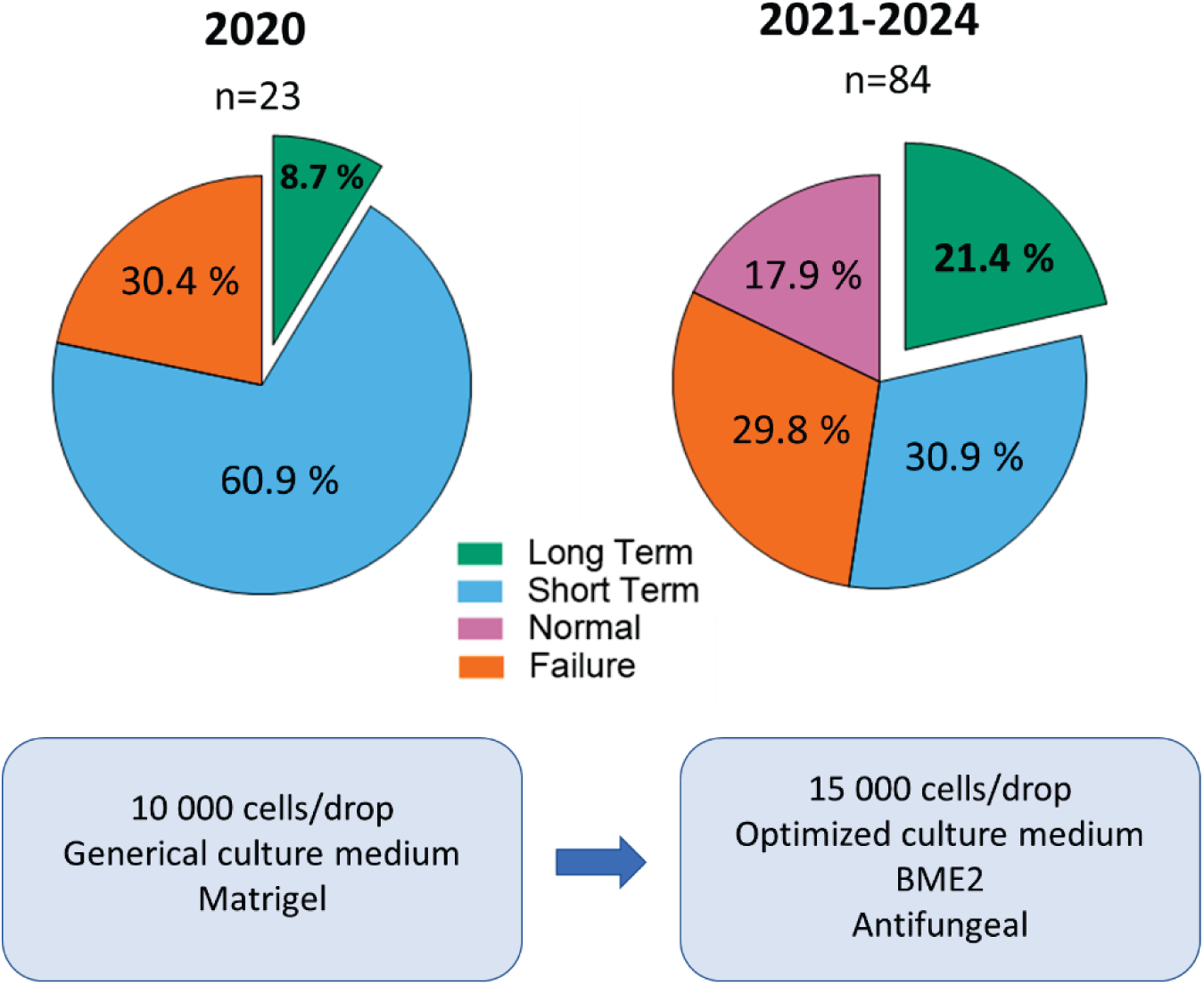
Evolution of the long-term PDTO establishment rate following optimization of culture conditions.

**Figure S3.**
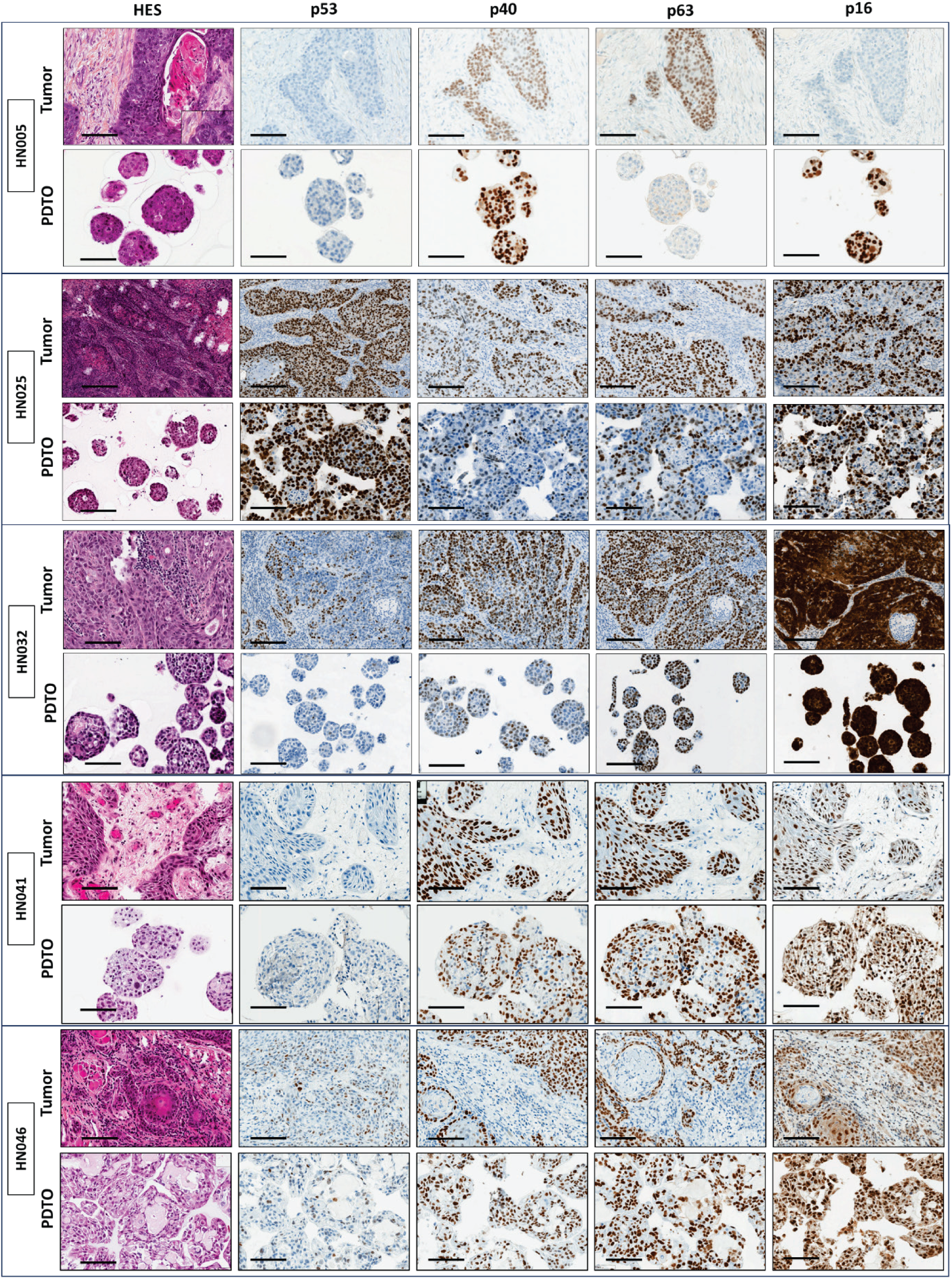
Histology and tumor marker expression in additional PDTO models. HES stain! ng and immunohistochemical analysis of HNSCC-associated biomarkers (pl6, p53, p40, and p63) in 21 PDTO lines and their paired tumors. Scale bar: 100 pm. (1/4)

**Figure S3.**
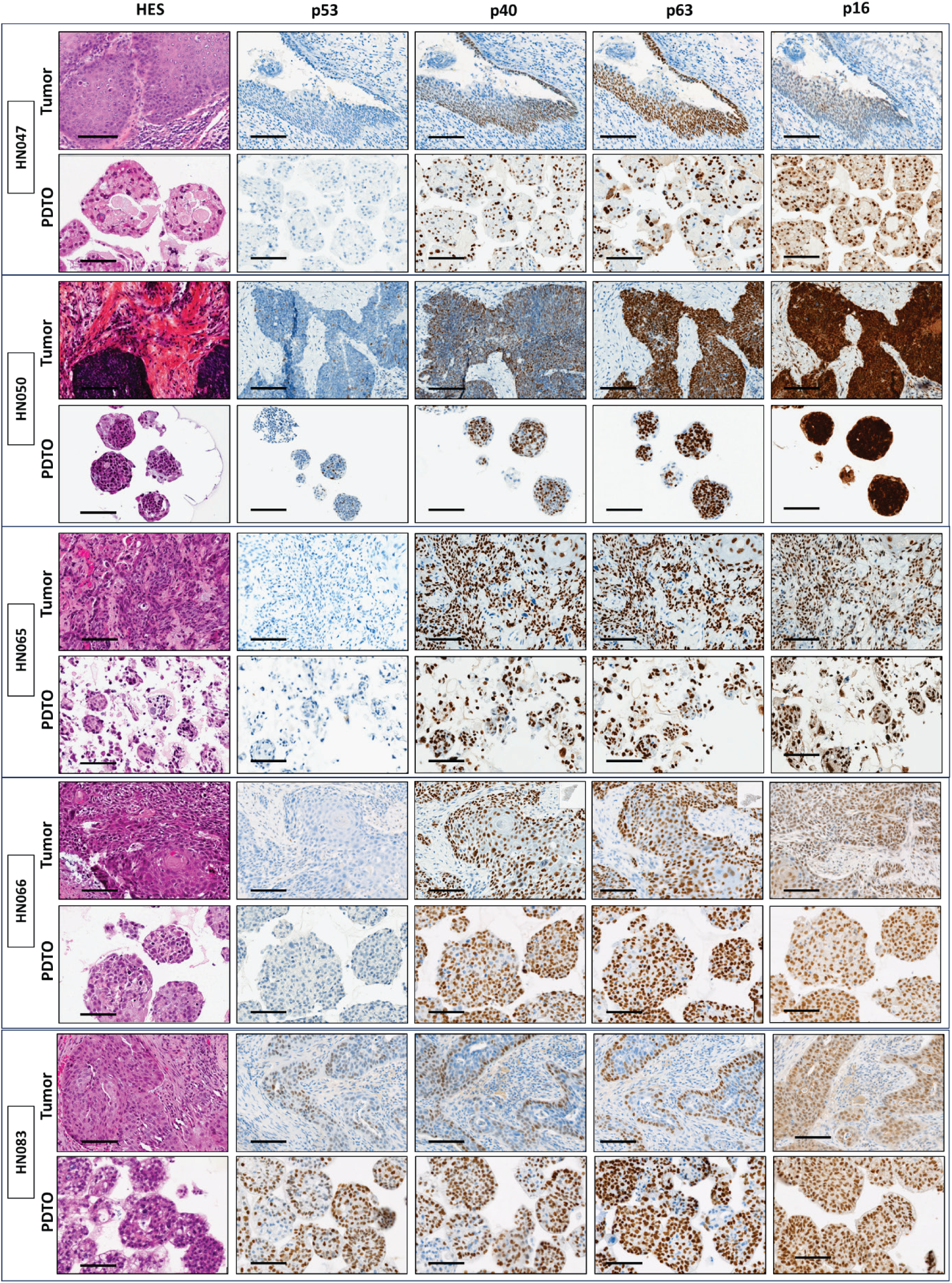
Histology and tumor marker expression in additional PDTO models. HES staining and immunohistochemical analysis of HNSCC-associated biomarkers (pl6, p53, p40, and p63) in 21 PDTO lines and their paired tumors. Scale bar: 100 pm. (2/4)

**Figure S3.**
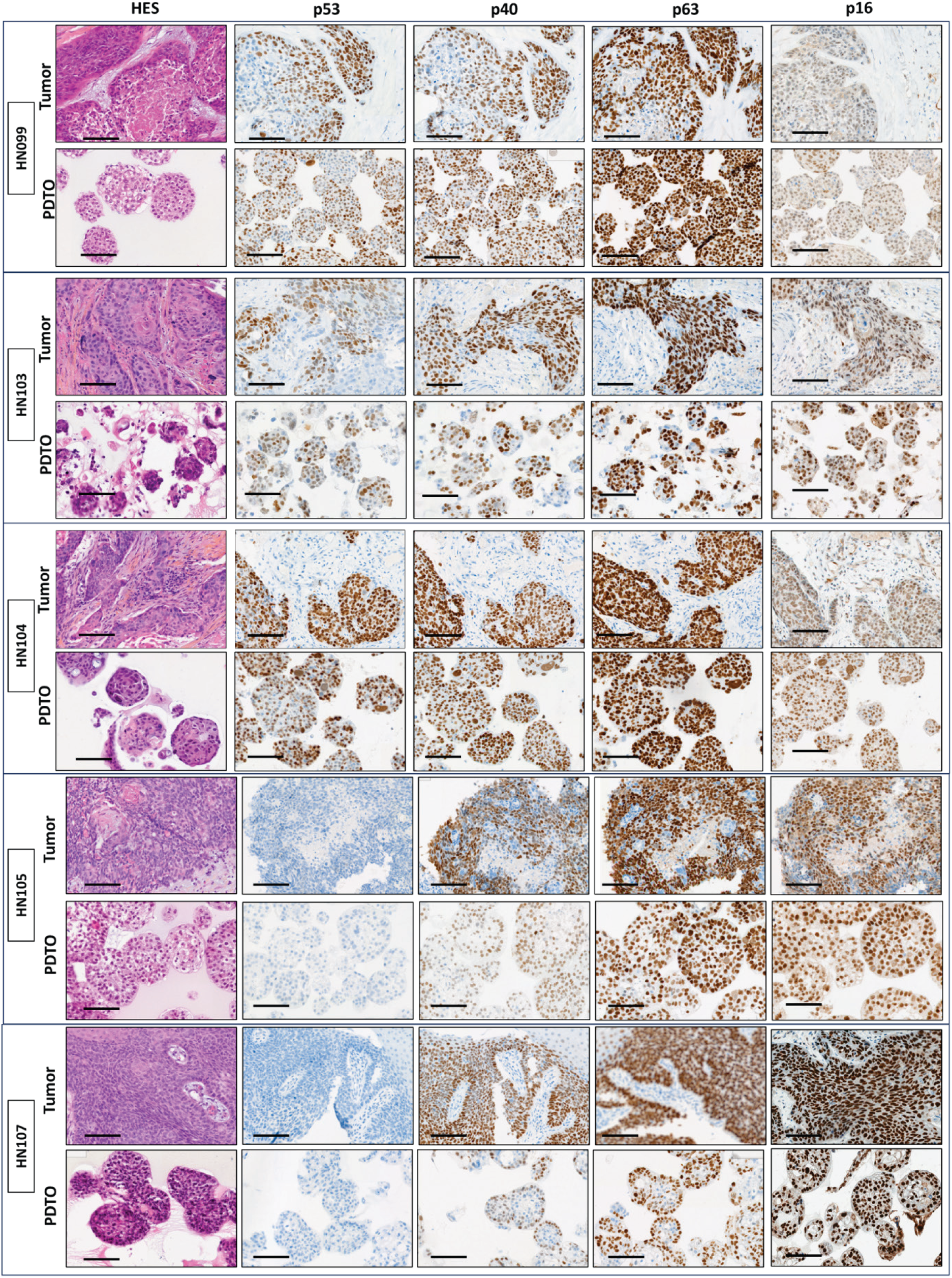
Histology and tumor marker expression in additional PDTO models. HES staining and immunohistochemical analysis of HNSCC- associated biomarkers (p!6, p53, p40, and p63) in 21 PDTO lines and their paired tumors. Scale bar: 100 pm. (3/4)

**Figure S3.**
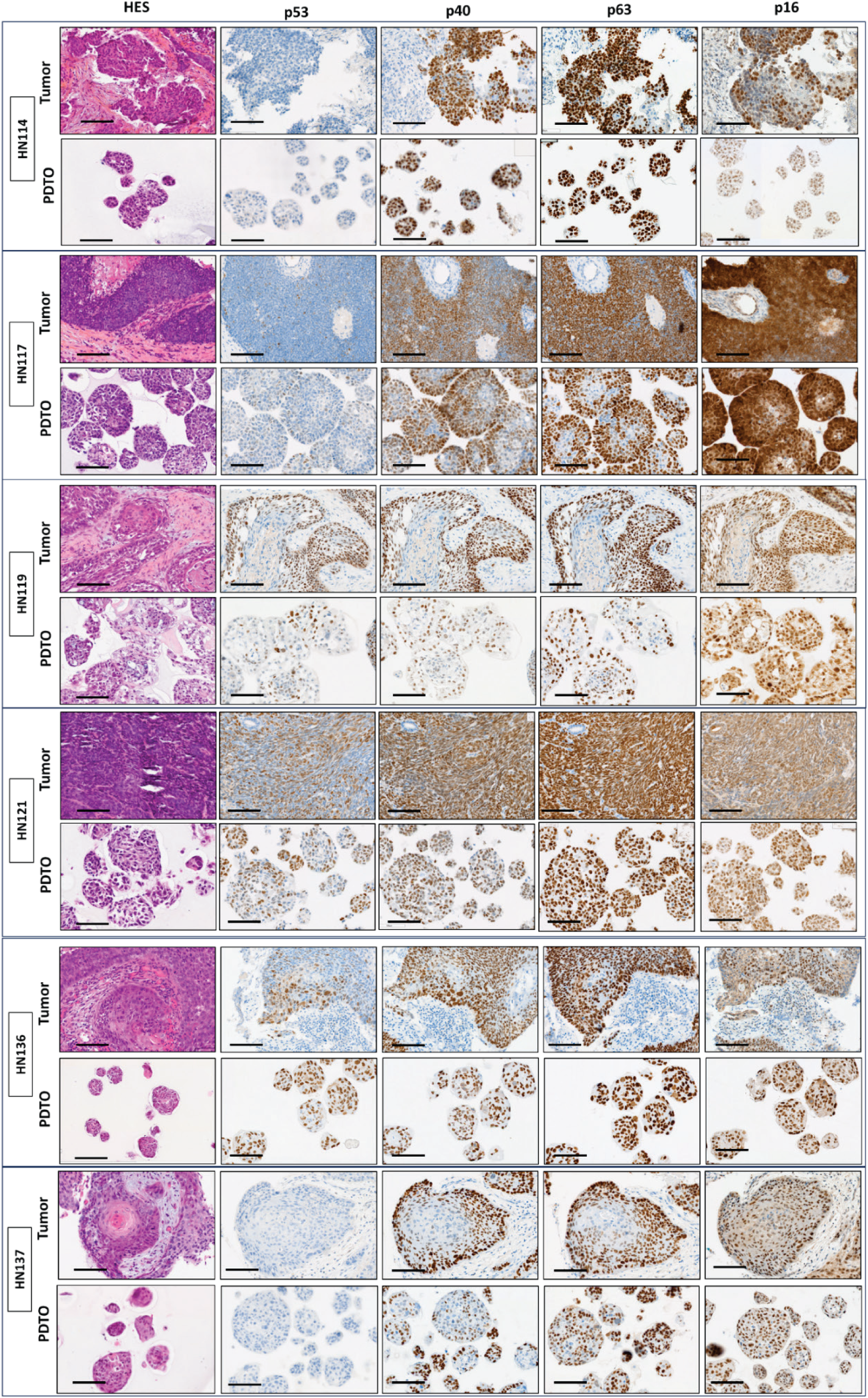
Histology and tumor marker expression in additional PDTO models. HES staining and immunohistochemical analysis of HNSCC-associated biomarkers (pl6, p53, p40, and p63) in 21 PDTO lines and their paired tumors. Scale bar: 100 pm. (4/4)

**Figure S4.**
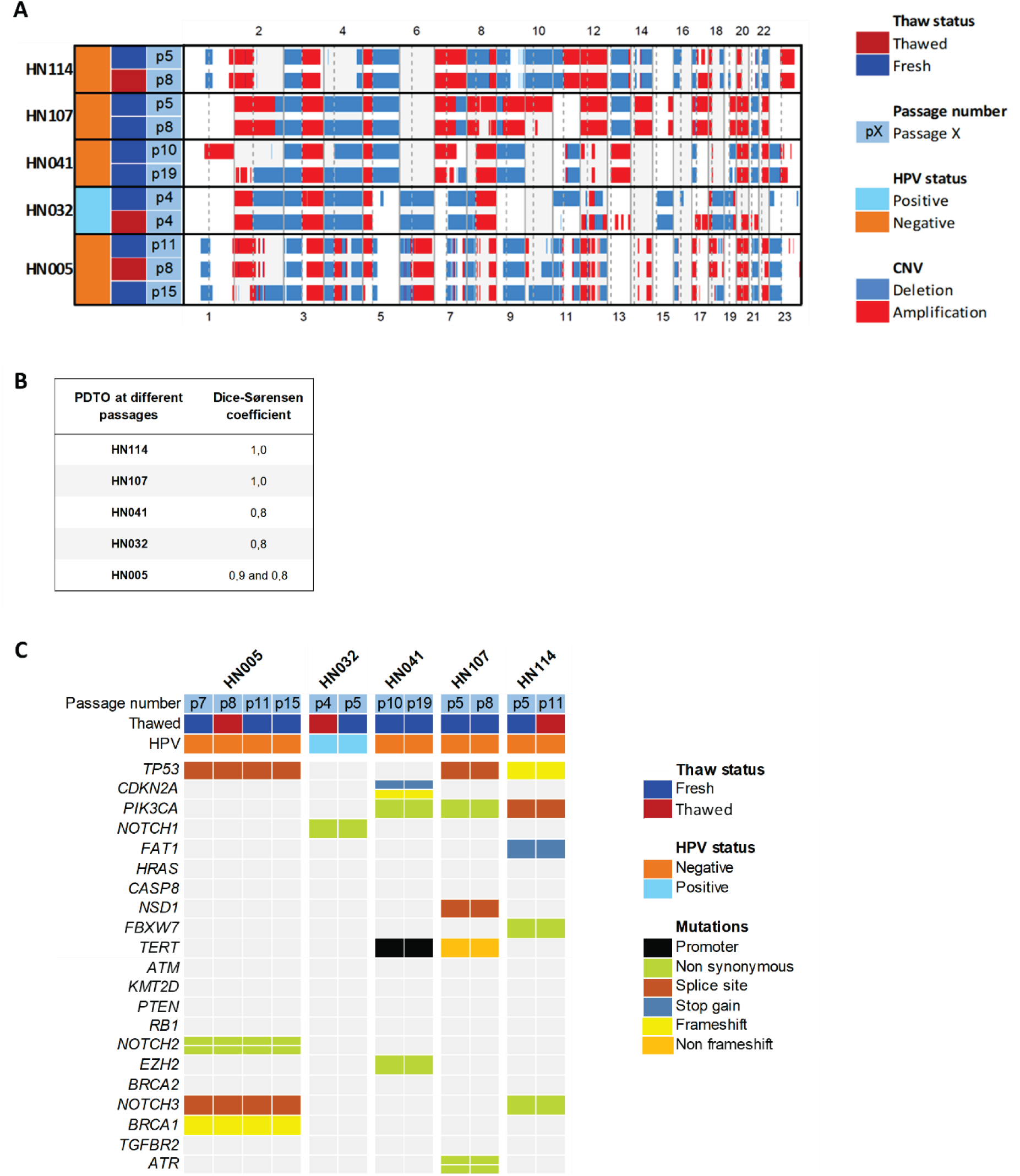
Genomic stability of PDTO across passages and after cryopreservation. **(A)** Copy-number variation (CNV) profiles analyzed by low-pass whole­genome sequencing (LP-WGS) in PDTO at different passages and after cryopreservation. **(B)** Correlation of CNV profiles across passages and after cryopreservation, calculated using the Dice-S0rensen coefficient. (C) Mutation status of 19 major HNSCC-associated genes across serial passages and after cryopreservation.

**Figure S5.**
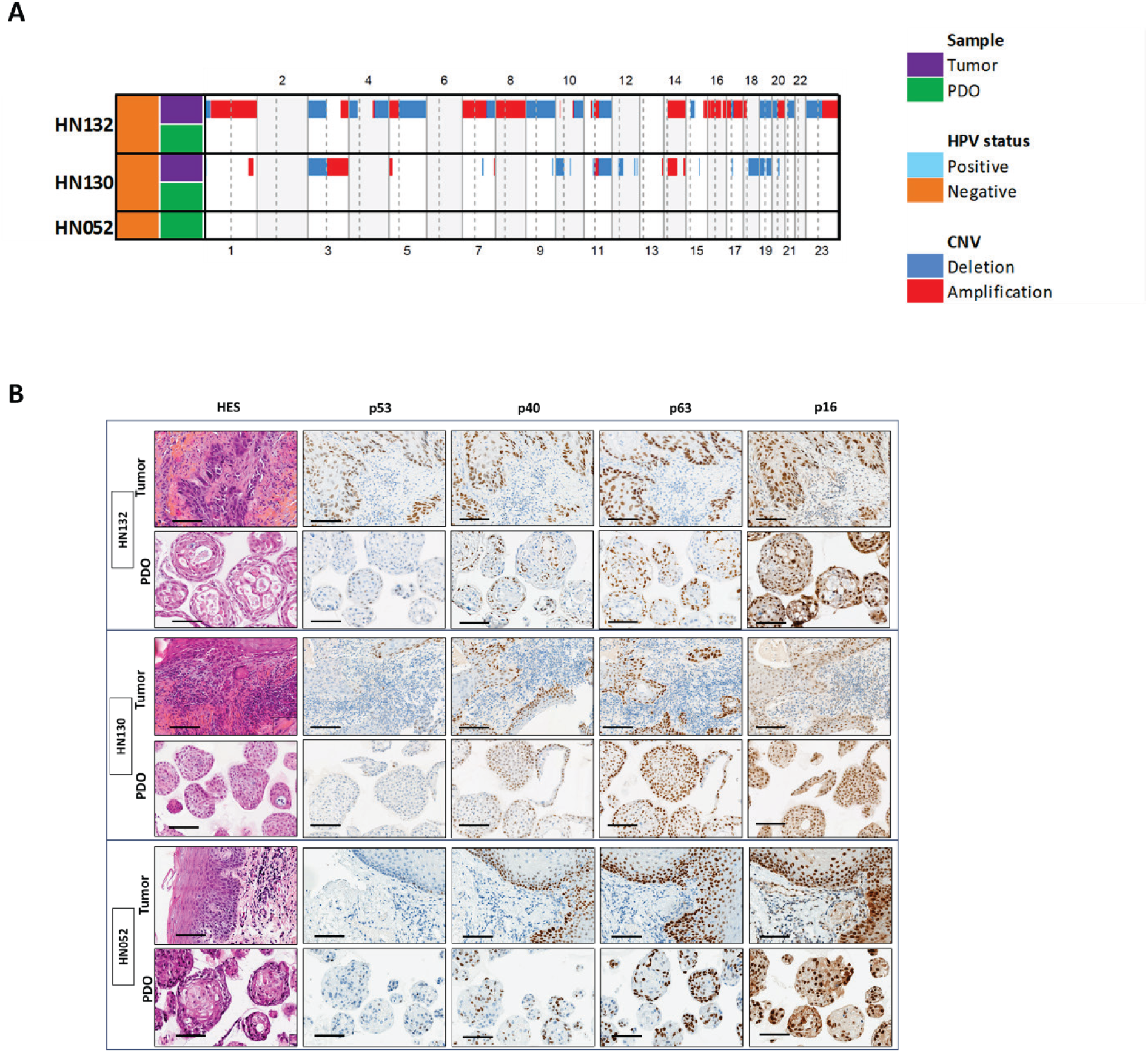
Characteristics of non-malignant PDO. **(A)** CNV profiles analyzed by LP-WGS in three PDO linesand their paired tumors. **(B)** HES staining and immunohistochemical analysis of p63 and Ki-67 in three PDO lines. Scale bar: 100 pm.

**Figure S6.**
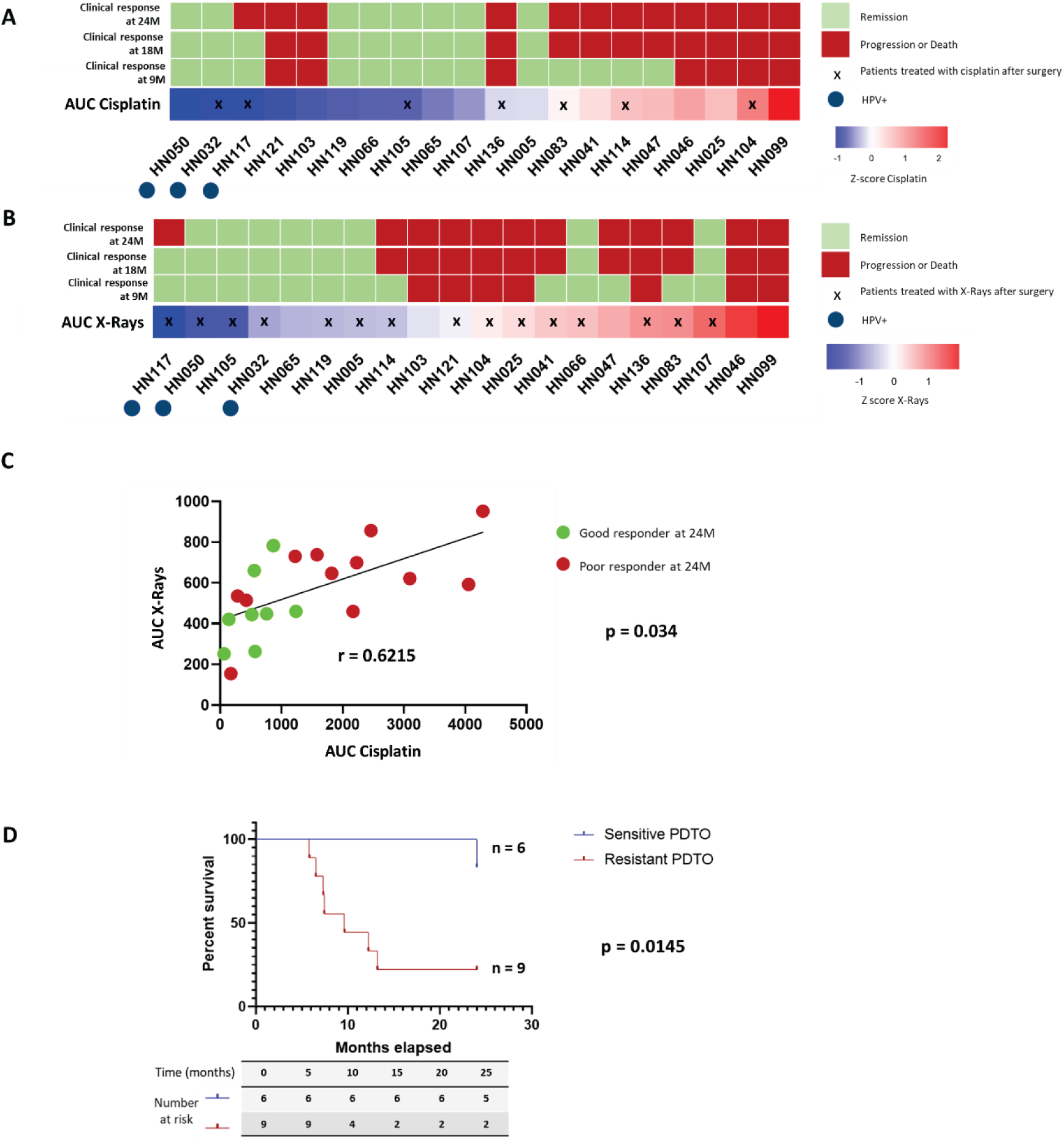
Extended analysis of the association between PDTO response and clinical outcome. Heatmaps ranking **(A)** cisplatin and **(B)** X-ray AUC values. Clinical response was defined as good {remission) or poor {relapse or death). Crosses indicate patients who received postoperative cisplatin and radiotherapy, **(C)** Correlation between cisplatin and X-ray AUC values. **(D)** Kaplan-Meier curves of DFS in patients who received postoperative radiotherapy stratified by PDTO sensitivity to X-rays.

**Table S1.**
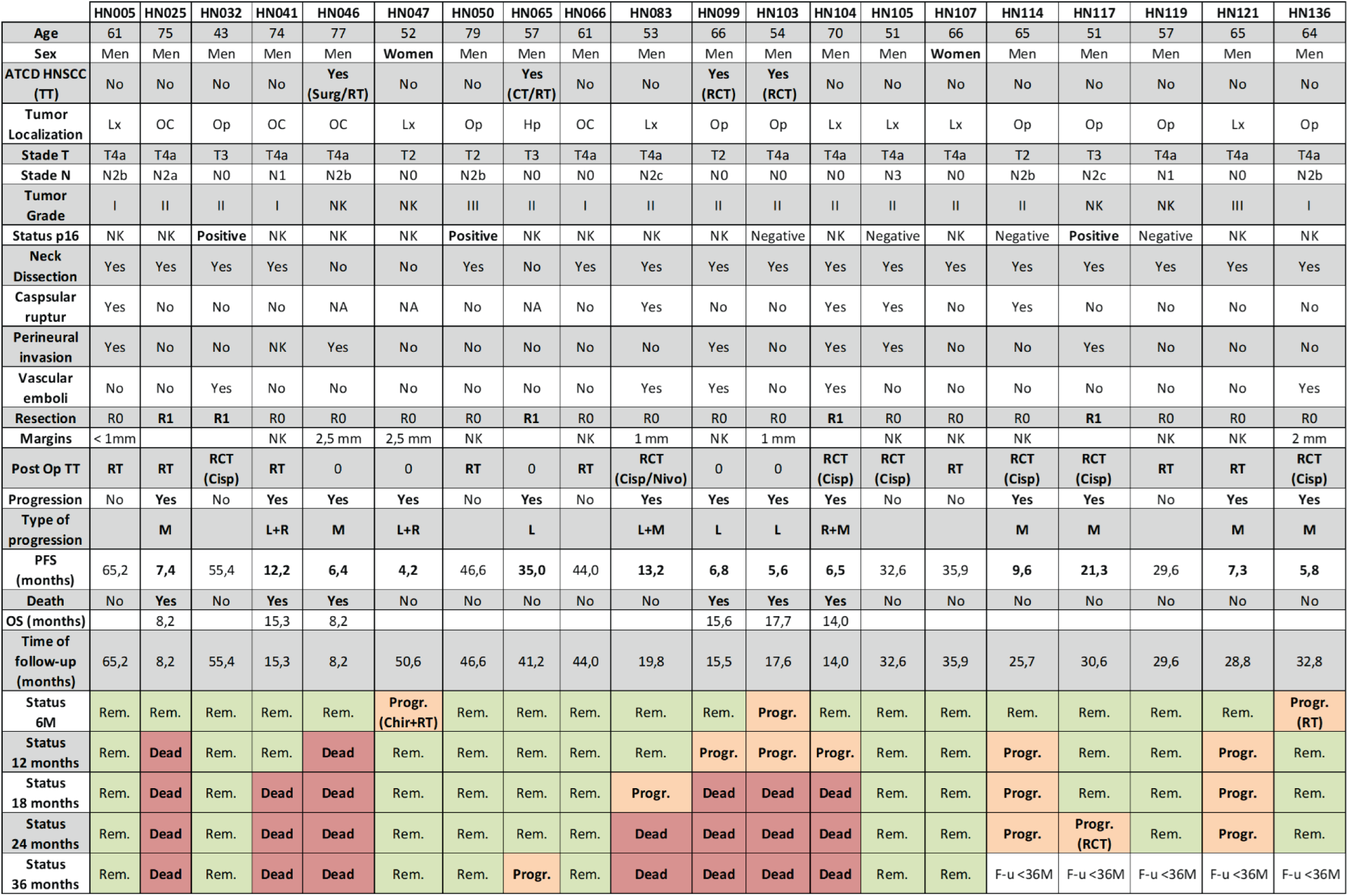
Clinical characteristics of the patients from whom the cohort of long-term PDTO were derived. Op: oropharynx; Lx: Larynx; Hp: hypopharynx; OC: oral cavity; TT: Treatment; RT: radiotherapy; CT: chemotherapy; RCT: radiochemotherapy; RO: clear margin; RI: positive margin; Cisp: cisplatin; Nivo: nivolumab; PFS: progression free survival; OS: overall survival; f/u: follow up; Prog.: progression; Rem.: remission; M: months; NK: not know.

**Table S2.**
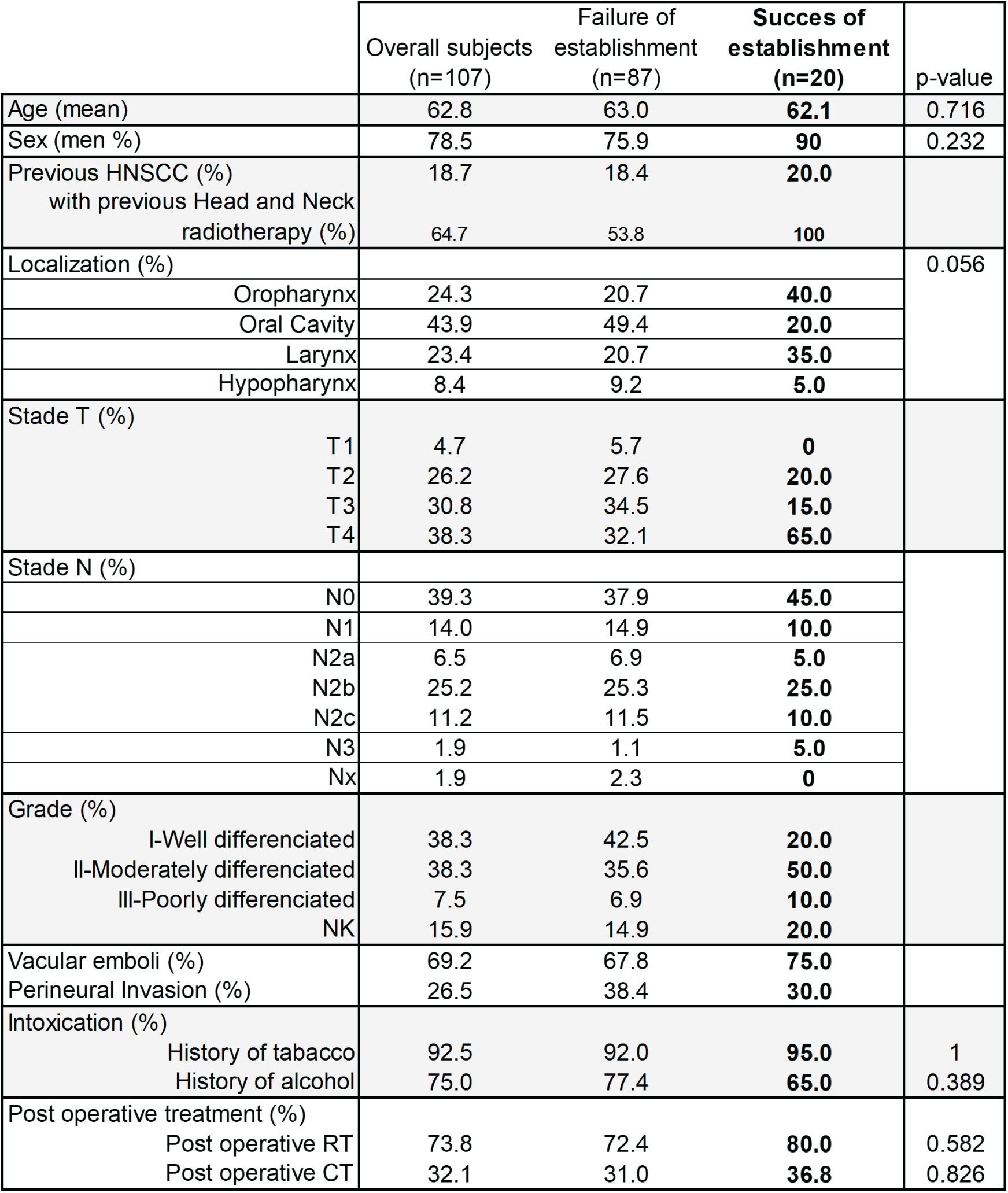
Clinical characteristics of the overall cohort and the long-term PDTO cohort. p-values comparing successful versus unsuccessful establishment are shown in the right column. RT: radiotherapy; CT: chemotherapy.

**Table S3.** Factors associated with successful establishment of long-term and short-term PDTO. Patients with missing data were excluded from the analysis. For short-term PDTO, normal organoids were excluded (n = 10). RT: radiotherapy; CT; chemotherapy.

|  | Long-term PDO |  |  |  | Short-term PDO |  |  |  |
| --- | --- | --- | --- | --- | --- | --- | --- | --- |
|  | N | OR | 95% IC | p value | N | OR | 95% IC | p value |
| Age | 107 | 0.99 | [0.94;1.04] | **0.7** | 93 | 0.97 | [0.93;1.02] | **0.2** |
| Age ≥ 68 years old | 107 | 0.78 | [0.23;2.26] | **0.7** | 93 | 0.65 | [0.26;1.62] | **0.3** |
| Sex | 107 |  |  |  | 93 |  |  |  |
| Men |  |  |  |  |  |  |  |  |
| Women |  | 0.35 | [0.05;1.35] | **0.2** |  | 0.52 | [0.20;1.40] | **0.2** |
| History of tobacco | 107 |  |  |  | 93 |  |  |  |
| No |  |  |  |  |  |  |  |  |
| Yes |  | 1.66 | [0.27;32.0] | **0.6** |  | 4.00 | [0.91;20.7] | **0.071** |
| History of alcohol | 104 |  |  |  | 90 |  |  |  |
| No |  |  |  |  |  |  |  |  |
| Yes |  | 0.54 | [0.19;1.62] | **0.3** |  | 1.48 | [0.54;3.94] | **0.4** |
| Previous HNSCC | 106 |  |  |  | 92 |  |  |  |
| No |  |  |  |  |  |  |  |  |
| Yes |  | 1.41 | [0.45;4.04] | **0.5** |  | 0.50 | [0.19;1.32] | **0.2** |
| Grade of differentiation | 107 |  |  |  | 93 |  |  |  |
| I |  |  |  |  |  |  |  |  |
| II |  | 2.98 | [0.90;11.7] | **0.087** |  | 1.88 | [0.68;5.40] | **0.2** |
| III |  | 3.08 | [0.37;20.0] | **0.2** |  | 1.95 | [0.38;14.7] | **0.5** |
| NK |  | 2.85 | [0.60;13.7] | **0.2** |  | 1.19 | [0.36;4.19] | **0.8** |
| Post-operative RT | 107 |  |  |  | 93 |  |  |  |
| No |  |  |  |  |  |  |  |  |
| Yes |  | 1.52 | [0.50;5.73] | **0.5** |  | 1.37 | [0.51;3.60] | **0.5** |
| Post-operative CT | 106 |  |  |  | 92 |  |  |  |
| No |  |  |  |  |  |  |  |  |
| Yes |  | 1.30 | [0.44;3.60] | **0.6** |  | 0.95 | [0.38;2.46] | **> 0.9** |
| Percentage of tumoral cells | 105 | 1.01 | [0.99;1.03] | **0.3** | 91 | 0.99 | [0.98;1.01] | **0.4** |

